# Analytical Choices Impact the Estimation of Rhythmic and Arrhythmic Components of Brain Activity

**DOI:** 10.1101/2025.09.24.678322

**Authors:** Jason da Silva Castanheira, Mathieu Landry, Stephen M. Fleming

## Abstract

Brain activity comprises both rhythmic (periodic) and arrhythmic (aperiodic) components. These signal elements vary across healthy aging, and disease, and may make distinct contributions to conscious perception. Despite pioneering techniques to parameterize rhythmic and arrhythmic neural components based on power spectra, the methodology for quantifying rhythmic activity remains in its infancy. Previous work has relied on parametric estimates of rhythmic power extracted from *specparam*, or estimates of rhythmic power obtained after detrending neural spectra. Variation in analytical choices for isolating brain rhythms from background arrhythmic activity makes interpreting findings across studies difficult. Whether these current approaches can accurately recover the independent contribution of these neural signal elements remains to be established. Here, using simulation and parameter recovery approaches, we show that power estimates obtained from detrended spectra conflate these two neurophysiological components, yielding spurious correlations between spectral model parameters. In contrast, modelled rhythmic power obtained from *specparam*, which detrends the power spectra and parametrizes brain rhythms, independently recovers the rhythmic and arrhythmic components in simulated neural time series, minimising spurious relationships. We validate these methods using resting-state recordings from a large cohort. Based on our findings, we recommend modelled rhythmic power estimates from specparam for the robust independent quantification of rhythmic and arrhythmic signal components for cognitive neuroscience.

## Introduction

Brain activity is composed of both rhythmic (periodic) and arrhythmic (aperiodic) signal components observed across spatiotemporal scales (Buzsáki & Draguhn, 2004; Buzsáki & Watson, 2012; Donoghue et al., 2020). Rhythmic signal components appear as peaks in the frequency domain that sit above the arrhythmic background (Buzsáki et al., 2013; Donoghue et al., 2020; Wen & Liu, 2016; L. E. Wilson et al., 2022, 2024). Quantifying rhythmic oscillations and relating these rhythms to behaviour has been a longstanding goal of neuroscientific research (Berger, 1929; Herrmann et al., 2016). Oscillations in cortical field potentials are amenable to computational modelling (Breakspear et al., 2010; H. R. Wilson & Cowan, 1972), are theorized to relate to the synchrony of populations of neurons (Baillet, 2017; Buzsáki et al., 2012; Wang, 2010), and are believed to organize brain activity temporally and spatially (Fries, 2005; Schölvinck et al., 2010; Singer, 2013; Varela et al., 2001). Over a century of research (Mushtaq et al., 2024) has shown that brain rhythms relate to perceptual and cognitive processes (Arnal & Giraud, 2012; Baillet, 2017; Samaha et al., 2020a), can differentiate individuals from one another (Castanheira et al., 2024; da Silva Castanheira et al., 2021; da Silva Castanheira, Poli, et al., 2024), and are altered by neurological and psychiatric diseases (Gallego-Rudolf et al., 2024; Heinrichs-Graham et al., 2014; Wiesman, Castanheira, et al., 2022; Wiesman et al., 2023).

Arrhythmic signal components, in contrast, are characterized by a power law (1/f^𝒳^) distribution in frequency space (Donoghue et al., 2020; Gao et al., 2017). The arrhythmic signal component has been traditionally treated as background noise, and something to be removed (Donoghue et al., 2020; Groppe et al., 2013). More recent investigations, however, challenge this viewpoint and suggest that the arrhythmic spectral component fluctuates alongside the demands of cognitive tasks (Cunningham et al., 2023; Gyurkovics et al., 2022; Waschke et al., 2021), reflects behavioural traits (Lu et al., 2024; B. D. Ostlund et al., 2021), accounts for the observation of flatter power spectra with increasing age (Voytek et al., 2015a; L. E. Wilson et al., 2022), and is altered by disease (da Silva Castanheira, Wiesman, et al., 2024; Donoghue, 2024; Wiesman et al., 2023). In addition, recent computational work and growing empirical evidence have led to the hypothesis that the arrhythmic exponent reflects a physiological balance between excitatory (E) and inhibitory (I) neural activity – a core property of brain dynamics that shapes neural computation, information flow, and network stability (Brake et al., 2024; Chini et al., 2022; Gao et al., 2017; Maschke et al., 2023; Weijs et al., 2025; Wiest et al., 2023).

A growing literature suggests that both arrhythmic and rhythmic neurophysiological activity correlate with behaviour and subjective experience (Koenig & He, 2025; Samaha et al., 2017; Wöstmann et al., 2019). Whether these neural signal components independently contribute to behaviours remains to be definitively demonstrated. To address such questions, methodological approaches to quantifying both rhythmic and arrhythmic signal components should be explored.

Despite a growing interest in understanding the behavioural and mechanistic consequences of arrhythmic brain activity, the methodology for distinguishing rhythmic from arrhythmic activity remains in its infancy. Previous work relied on measures of relative power, which do not account for the power law (1/f^𝒳^) background activity, muddying the relationship between rhythmic and arrhythmic components (Davidson et al., 2022; Rempe et al., 2023; Samaha et al., 2022). Other researchers advocate for spectral detrending approaches which rely on computational modelling, such as *specparam*, which decomposes power spectra into their constituent signal elements (Donoghue et al., 2020; Wen & Liu, 2016; L. E. Wilson et al., 2022, 2024). The *specparam* model and its derivatives assume that neural power spectra are composed of independent contributions from rhythmic and arrhythmic activity, which are in turn associated with unique model parameters (i.e., the 1/f slope and Gaussian peaks)

(Donoghue et al., 2020; Medrano et al., 2025; L. E. Wilson et al., 2022, 2024). The *specparam* algorithm iteratively searches for Gaussian rhythmic peaks that sit above the power law (1/f^𝒳^), and parametrizes their centre frequency, amplitude, and bandwidth. The height of these Gaussian peaks (i.e., amplitude) reflects the total power of a given brain rhythm (Figure 1b). In addition, the arrhythmic signal component is characterised by two parameters: the offset and slope (i.e., exponent; 𝒳). The arrhythmic exponent represents the slope of the 1/f-like function decay, with low values indicating flatter spectra. Note that ‘arrhythmic exponent’ and ‘spectral slope’ are used interchangeably in the literature, with the spectral slope representing the signed exponent. The arrhythmic slope may also be parametrized with an optional spectral ‘knee’ parameter, which models spectral bends often observed in intracranial data and spectra across a large frequency range. These signal elements are summed together in the frequency domain to generate the broadband signal observed in neurophysiological recordings (Donoghue et al., 2020; L. E. Wilson et al., 2024).

**Figure 1:**
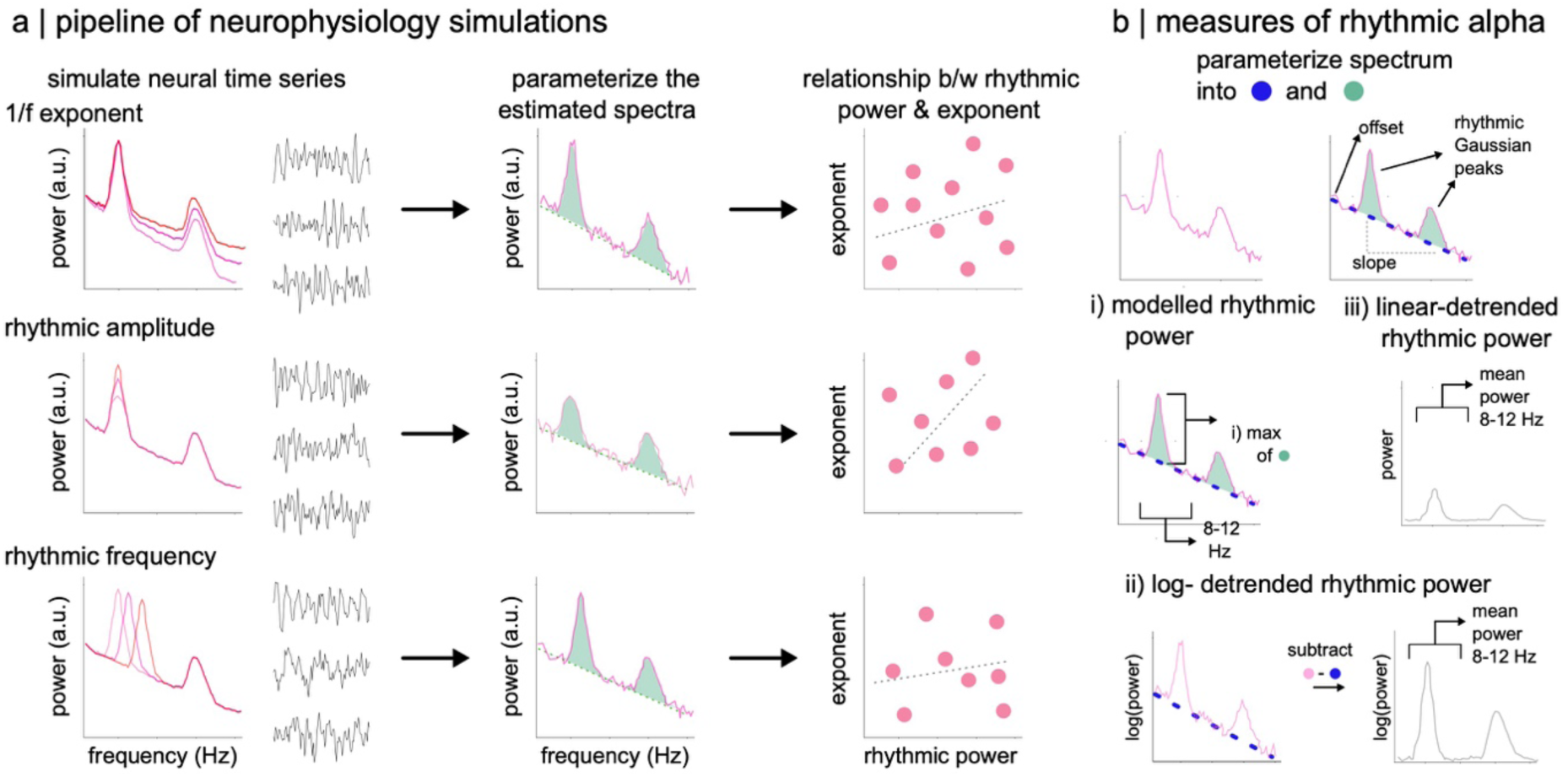
Schematic of simulations & definition of rhythmic power. (a) Schematic of the pipeline of neural time series simulations. Neural time series data were simulated based on parameters of the *specparam* model in frequency space. These parameters included the slope of the 1/f arrhythmic activity (top panel), the amplitude of rhythmic activity (middle panel) or the center frequency of rhythmic activity (bottom panel). We simulated neural time series with various ground truth relationships between the arrhythmic and rhythmic components. These simulated neural time series were transformed into the frequency domain and parameterized. The relationship between the simulated components was then evaluated based on the parameterized outputs. (b) Schematic of different methodological choices for estimating rhythmic neural power. The rhythmic component of brain activity can be defined in three ways: i) as the maximum amplitude of modelled Gaussians within a defined frequency range (e.g., the alpha range 8-12 Hz). We define this method as modelled rhythmic power. Rhythmic power can also be computed as detrended rhythmic power obtained by subtracting the modelled 1/f arrhythmic component from the power spectrum and computing the mean residual variance within a predefined narrow-band range. This subtraction can be done in either in ii) log or iii) linear space.

**Figure 2:**
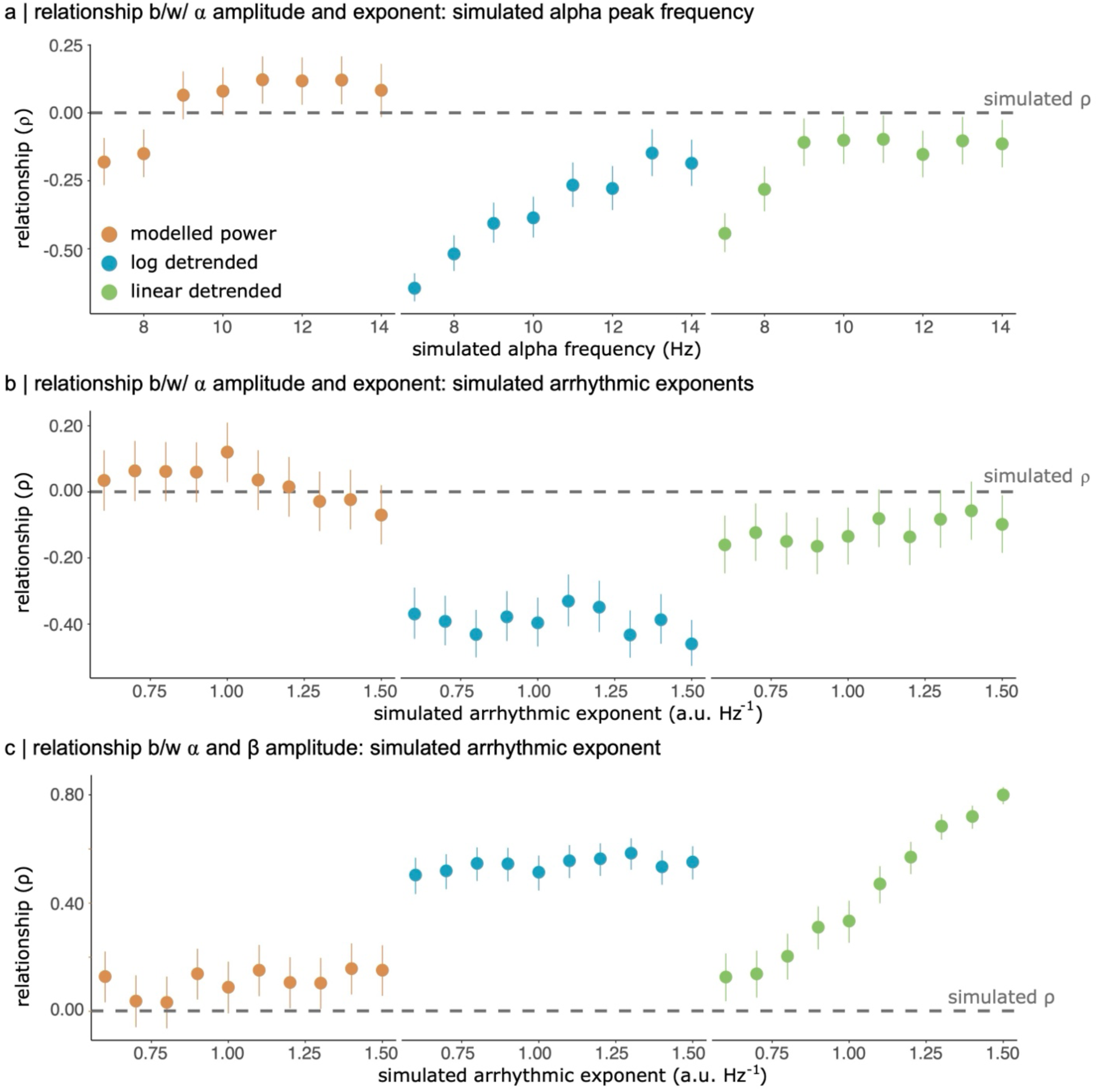
Spectral detrending induces spurious correlations between spectral model outputs. (a) Estimated correlation between rhythmic alpha power and arrhythmic exponent at various simulated alpha center frequencies. The ground truth correlation is zero, indicated by the dotted line. Modelled power computed as the maximum amplitude of modelled Gaussian peaks is closer to the ground truth relationship between simulated alpha power and exponent (i.e., ⍴ =0.00). Spurious negative correlations of large magnitude (⍴ < -0.40) between alpha power and arrhythmic exponent were obtained for log-detrended power. This is especially evident when the center frequency of the alpha oscillation lies at the edge of the narrow-band definition (i.e., 7-8 Hz). (b) Estimated correlation between rhythmic alpha power and arrhythmic exponent for various levels of simulated arrhythmic exponents. Modelled alpha power recovers the ground truth relationship significantly better than detrending approaches. Log-detrended power induced large spurious correlations between alpha power and arrhythmic exponent for all simulated values of the arrhythmic exponent. (c) Scatter plot of the relationship between rhythmic alpha and beta power for various simulated arrhythmic exponents. While synthetic time series data were simulated to have no linear relationship between alpha and beta amplitudes, spurious correlations were obtained between alpha and beta band power for log- and linear-detrending methods. In contrast, modelled power (left-most panel) shows no such spurious relationship between alpha and beta power, accurately recovering the ground truth null correlation between these parameters. Error bars represent 95% CI.

**Figure 3:**
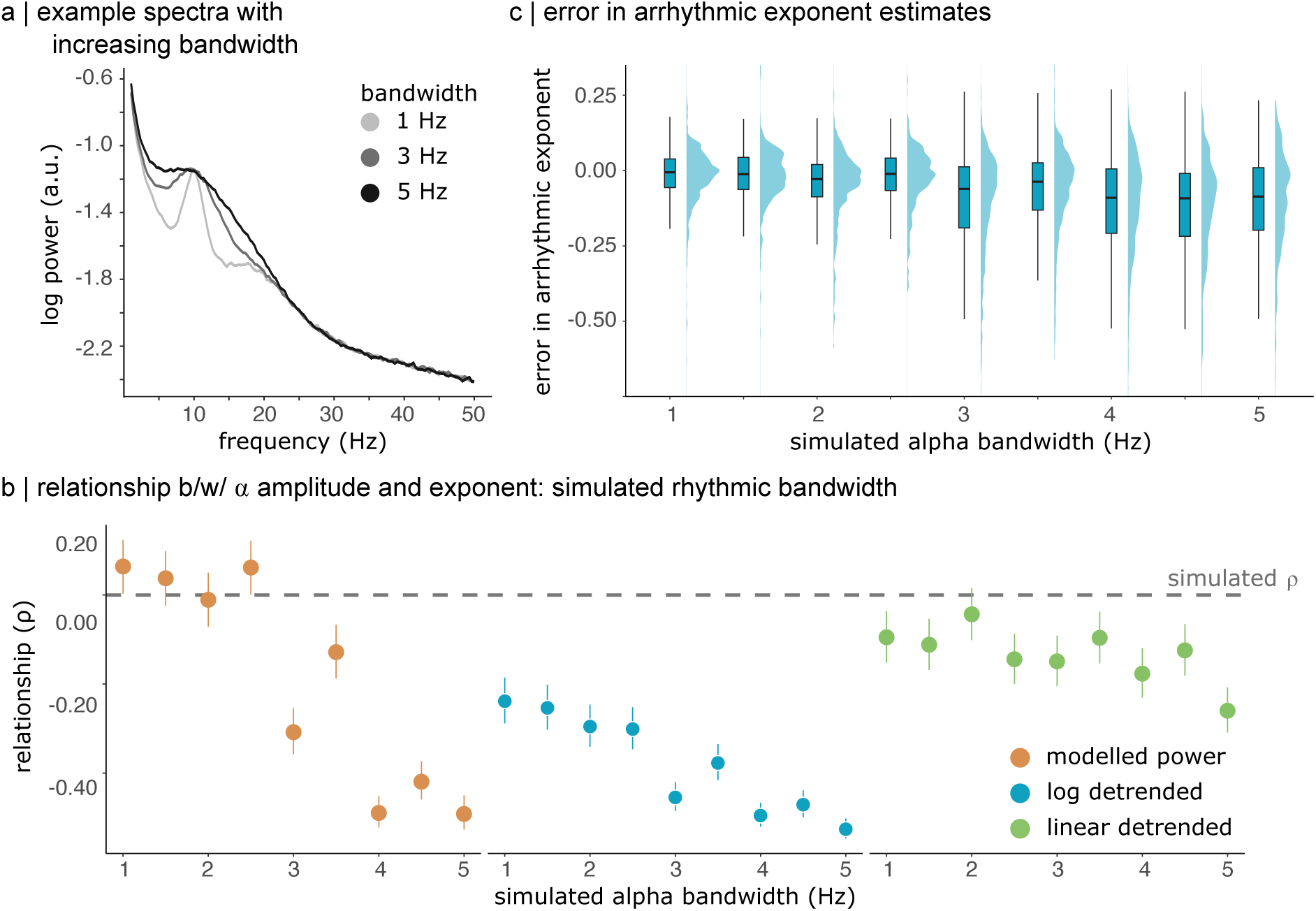
Broad spectral peaks challenge the independent recovery of spectral model outputs. (a) Average power spectra across simulations for alpha peak bandwidths of 1, 3, and 5 Hz. Larger alpha peak bandwidths yield broad peaks in the spectral domain that sit above the 1/f arrhythmic background. (b) Estimated correlation between rhythmic alpha power and arrhythmic exponent for various levels of simulated alpha peak bandwidth. Linear detrended alpha power recovers the ground truth relationship better than the other two methodologies on average. Log-detrended power induced large spurious correlations between alpha power and arrhythmic exponent for all simulated values of the arrhythmic exponent. Modelled rhythmic power recovers the ground truth relationship accurately for narrow spectral peaks (bandwidths < 3). Error bars represent 95% CI. (c) Distributions of error in the estimated arrhythmic exponent across the various values of simulated alpha bandwidths. Errors in the recovered arrhythmic exponent are more variable for larger simulated spectral bandwidths (>3Hz). This suggests that the *specparam* model struggles to fit broad spectral peaks, impacting the estimation of the arrhythmic component at this specific SNR level.

**Figure 4:**
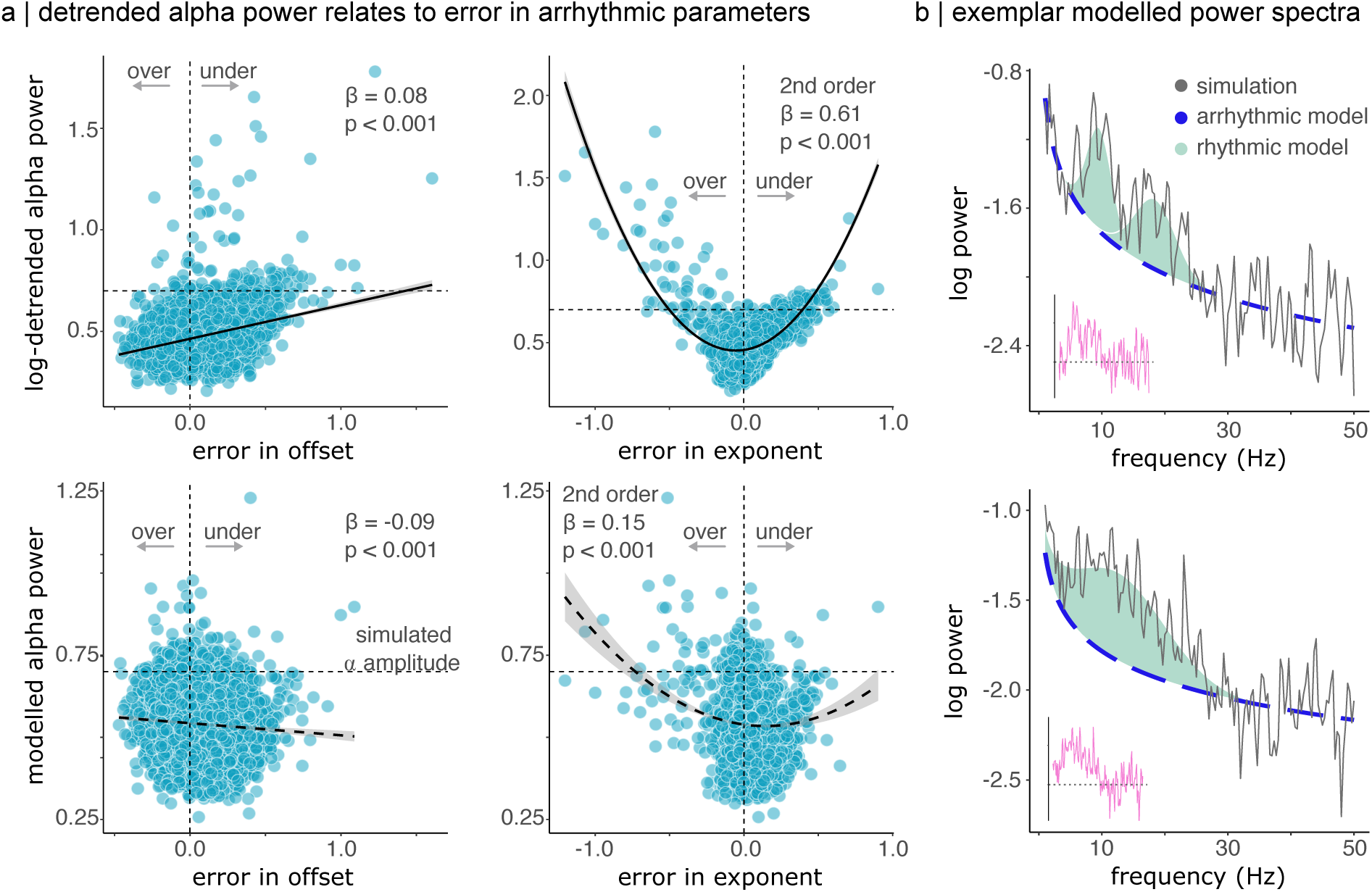
Error in arrhythmic parameter estimates predicts corrected but not modelled alpha power. (a) Scatter plot of the relationship between alpha power (top panel, log-detrended power; bottom panel, modelled power) and error in the estimate of arrhythmic parameters. Log-detrended alpha power is linearly related to error in the arrhythmic offset, with larger estimates of alpha power observed for underestimated values in arrhythmic offset. Error in arrhythmic exponent nonlinearly related to alpha power, with extreme under- and over-estimated values of the exponent predicting large alpha power. Dashed lines indicated the simulated (ground truth) alpha power, and zero error along the y- and x- axis respectively. Modelled alpha power does not meaningfully relate to error in arrhythmic parameters. (b) Example neural power spectra. The top panel depicts an accurately modelled power spectrum, while the bottom panel illustrates a poorly parameterized power spectrum. Black lines indicate the simulated spectrum, the blue dashed line the arrhythmic model, and the inlaid pink line the detrended alpha power. Modelled rhythmic Gaussians sit atop the modelled arrhythmic slope—together they compose the *specparam* model. In the bottom panel, the modelled arrhythmic offset is estimated to be lower than the ground-truth simulated value.

Even within model-based approaches, methodological choices differ: some papers use the Gaussian height parameters from the *specparam* model as a measure of rhythmic power (Donoghue et al., 2020; Hill et al., 2022; B. Ostlund et al., 2022; Wiest et al., 2023), whereas others quantify the residual variance after subtracting the arrhythmic component from the neural power spectrum, either in log-log or linear space (i.e., detrending analyses) (da Silva Castanheira, Wiesman, et al., 2024; Lu et al., 2024; Merkin et al., 2022, 2023; Wiesman et al., 2023). Recent findings even question data-driven detrending approaches altogether (Brake et al., 2024).

Brake et al. find that the arrhythmic background activity can be altered either in a multiplicative or additive manner, which impacts the kind of spectral detrending approach which should be performed (i.e., detrending in log or linear space). As changes in the arrhythmic component cannot be determined as multiplicative or additive in advance, the choice of detrending method may be inappropriate. In contrast to the recommendations of the *specparam* algorithm, Brake et al. suggest that spectral detrending should be avoided entirely when clear oscillatory peaks are present (Brake et al., 2024). They suggest that unless clear physiological and biophysical justifications are provided, spectral detrending should not be performed. However, the analyses of Brake et al. do not explicitly model the rhythmic spectral peaks, making it inappropriate for our purposes of separating rhythmic and arrhythmic components.

More generally, the impact of these methodological choices on the decomposition of rhythmic and arrhythmic activity, and how they affect the conclusions drawn about links to behaviour, remains unknown. Without clear, evidence-based guidelines on how to appropriately decompose brain rhythms, the robustness, reproducibility, and interpretability of findings in the field remain underspecified.

Here we sought to explore how methodological choices in quantifying rhythmic activity from neural power spectra impact the interpretability of findings – first in simulation, and then in resting-state MEG recordings. Relying on simulated neural time series, we tested whether the estimated correlation between spectral model parameters recovered the ground truth for various approaches to distinguishing rhythmic and arrhythmic components. Our goal was to verify the robustness of various methods for quantifying rhythmic brain activity and demonstrate how analytical choices can yield diverging interpretations of empirical data.

## Methods

### Measures of rhythmic power

Rhythmic brain activity within a narrow frequency band (e.g. alpha 8-12 Hz) was estimated as either i) modelled rhythmic power, or detrended power in ii) log-log space and iii) linear space averaged within a narrow band (see Figure 1b). Modelled power consisted of the height of the Gaussian peak fit by the *specparam* algorithm within a pre-defined frequency range. In the case of multiple fitted Gaussian peaks, we retained the Gaussian peak of the highest amplitude. If *specparam* fit no Gaussian peak within the frequency band of interest, we assigned the amplitude a value of NaN or 0 (see <u>Simulations of neural time series</u> for further exploration of how to deal with missing values). For detrended spectral power, we relied on previous definitions of rhythmic activity as the residual variance in the observed power spectrum after subtracting the modelled 1/f arrhythmic component. Note that the subtraction between the observed spectrum and arrhythmic power can be performed in either linear or log-log space. This is equivalent to subtracting or dividing the arrhythmic component from the spectrum, respectively (Brake et al., 2024). We defined rhythmic power as the mean power of this residual variance within a pre-defined frequency range (e.g., 8-12 Hz for the alpha band; Figure 1b, bottom panel). We similarly explored the maximum power in the alpha band (8-12 Hz) in log and linear detrended spectra. Averaging within a narrowband or using the maximum power resulted in similar findings.

Note that we did not explore measures of ‘raw’ spectral power as suggested by Brake et al. as this method does not take any steps towards separating out the rhythmic and arrhythmic components of the power spectrum, and by definition, estimates of ‘raw’ power should be strongly correlated with the arrhythmic exponent.

### Simulations of neural time series

We generated over 20,000 simulated neural time series with a range of arrhythmic and rhythmic parameters. Neural time series were synthesized using the NeuroDSP toolbox (Cole et al., 2019), lasted 30 seconds, and had a sampling rate of 500 Hz. The choice of a brief 30-second recording was informed by previous empirical research, which established that this duration yields stable estimates of the power spectrum (da Silva Castanheira et al., 2021; Wiesman, da Silva Castanheira, et al., 2022). The NeuroDSP toolbox (Cole et al., 2019) simulates neural time series with given spectral properties. It separately simulates aperiodic and periodic neural time series, which are then summed together. For the arrhythmic component, the toolbox rotates white noise in frequency space to obtain the desired arrhythmic slope. For the periodic component, the desired time series is computed by constructing an array of amplitude coefficients that reflect all of the simulated spectral peaks. These coefficients are used to compute a summed series of cosine signals with random phase shifts. The resulting aperiodic and periodic time series are then summed together to generate the desired time series with the intended spectral properties.

The range of rhythmic and arrhythmic parameters used to generate the synthetic data was based on those extracted from the CamCAN dataset and previous computational work (see <u>Empirical dataset</u> for details) (L. E. Wilson et al., 2024).

Figure 1a depicts a schematic of the analysis pipelines applied to the simulated neural time series. First, to explore whether methodological choices in quantifying rhythmic activity lead to spurious relationships between the different parameters of the *specparam* model, we ran three sets of simulations. We manipulated either i) arhythmic exponent, ii) alpha amplitude, iii) alpha centre frequency, or iv) alpha peak bandwidth while holding all other parameter values constant. Within the simulated time series, there was no systematic relationship between arrhythmic exponent and rhythmic alpha power. Any significant correlation between these estimated components would therefore reflect a systematic bias introduced by the method chosen to quantify rhythmic activity.

We simulated 500 unique time series for each alpha amplitude, which ranged from 0.3 to 2.0 in steps of 0.2, with the alpha centre frequency and bandwidth held constant at 10 Hz and 2 Hz, respectively. We set the arrhythmic exponent to 1.0. Next, we simulated 500 unique time series for alpha centre frequencies ranging from 7 to 14 Hz in steps of 1 Hz. Alpha amplitude and arrhythmic exponent were held constant at 0.7 and 1.0, respectively. We simulated 500 unique time series for each value of the arrhythmic exponent, which varied from 0.6 to 1.5 Hz^-1^ in steps of 0.1 Hz^-1^. Alpha centre frequency, amplitude, and bandwidth were held constant for these simulations. For all of these simulations, we included a constant beta peak of 19 Hz, with an amplitude of 0.4 a.u. and a bandwidth of 5 Hz. Finally, we simulated 500 unique time series for each value of alpha bandwidth, which varied between 1 and 5 Hz in steps of 0.5 Hz. For these simulations, the value of alpha amplitude, peak frequency, and arrhythmic exponent were held constant.

Second, we tested whether the method for quantifying rhythmic brain activity impacts the recovery of a monotonic relationship between arrhythmic exponent and rhythmic alpha activity. To do so, we simulated 100 neural time series for 10 different effect sizes representing the relationship between alpha amplitude and arrhythmic exponent, starting from ⍴= 0.0 up to ⍴= 0.9 in steps of 0.1. Alpha amplitude ranged from 0.1 to 1.1 a.u., and arrhythmic exponent ranged from 0.5 to 1.5 Hz^-1^. Beta centre frequency, amplitude, and bandwidth were held constant at 19 Hz, 0.4 a.u. and 5Hz, while alpha centre frequency and bandwidth were held constant at 10 Hz and 2 Hz, respectively. We tested whether the recovered correlation between spectral parameters matched the simulated ground truth for the three methodologies of computing rhythmic alpha power—i.e., modelled power, log- and linear- detrending.

In a complementary analysis, we assessed the impact of missing values on the recovered relationships between alpha power and arrhythmic slope by either excluding data or replacing missing data with zero amplitude peaks. When defining rhythmic power as the height of the modelled Gaussian peak of *specparam*, users must decide whether to ignore spectra that do not contain a modelled peak within a pre-defined narrow band range of interest (e.g., the alpha band) or whether to replace missing values with zero (i.e., reflecting a Gaussian peak of height 0 above the arrhythmic background activity). The results of these analyses are presented in Figure 5.

**Figure 5:**
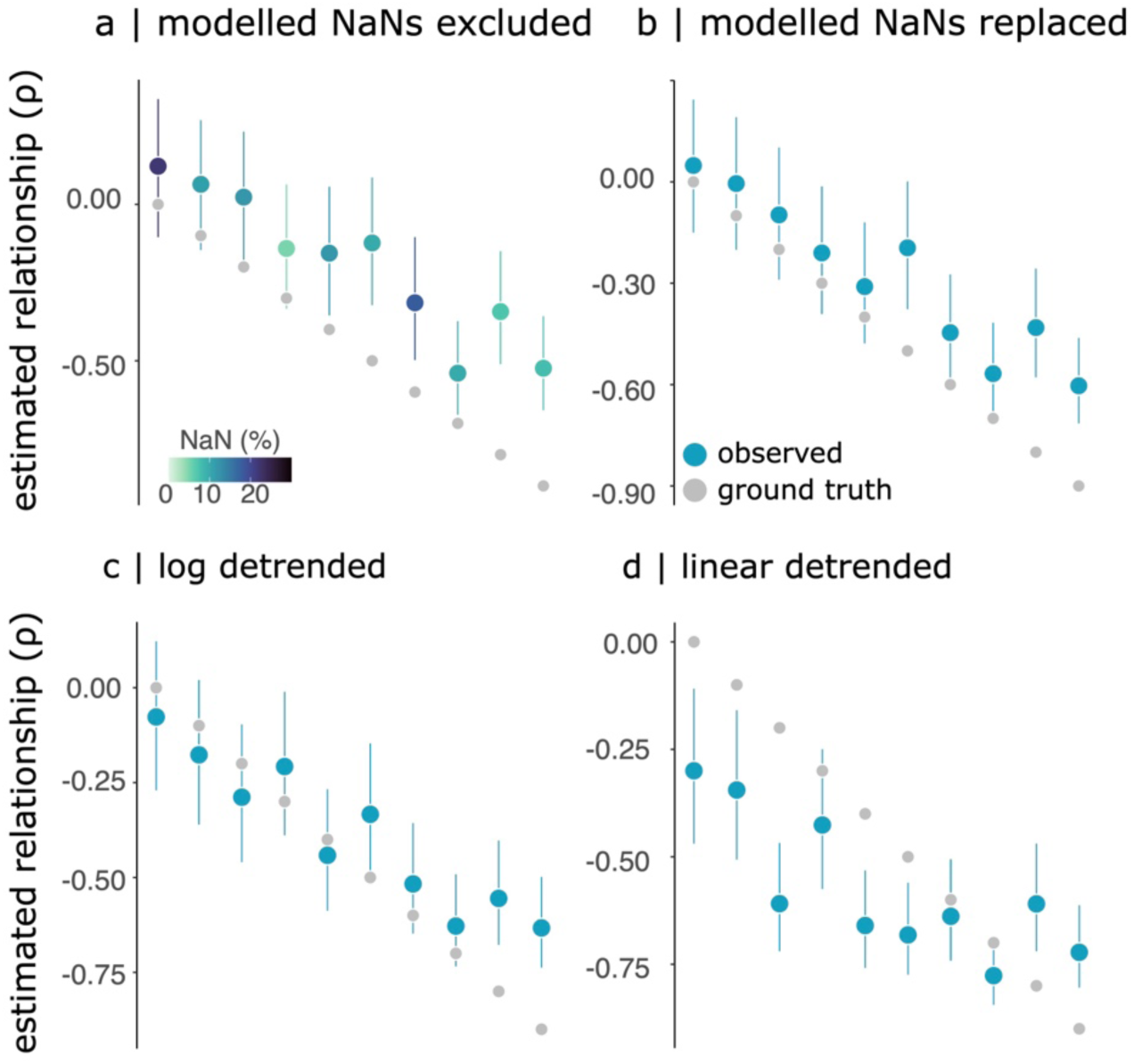
Replacing missing values with zeroes recovers simulated relationships. Simulated relationships between rhythmic alpha power and arrhythmic exponent for various effect sizes across the three methods of extracting alpha power. Ground truth values (grey dots) represent the simulated relationship between the two parameters used to generate neural time series data. Blue dots represent the estimated relationship between rhythmic alpha power and arrhythmic exponent for modelled power. Modelled power recovers the simulated relationship between alpha power and arrhythmic exponent reasonably well, except at the largest effect sizes (a). Modelled alpha power generates missing data when no Gaussian peak is fit within the alpha range. The colour of the dots represents the percentage of missing data for each simulation. The estimated relationships are closer to the ground truth when missing values of spectral peak amplitudes are replaced with zeroes (b) in comparison to excluding these values (a). Log-detrended alpha power similarly recovers the simulated relationships well (c). However, linear-detrended power systematically overestimates the strength of the linear relationship (d). Error bars represent 95% CI.

The power spectra of each simulated time series were computed using Welch’s method, utilizing 3-second windows with a 50% overlap as implemented in Brainstorm (Tadel et al., 2011).

### Parameterization of neural power spectra

To quantify the contribution of arrhythmic and rhythmic components to both empirical and synthetic time series, we parameterized power spectra using the *ms-specparam* tool in Brainstorm (Donoghue et al., 2020; Tadel et al., 2011; L. E. Wilson et al., 2024). We parameterized power spectra between 1 and 50 Hz for the simulated data and between 1 and 40 Hz for the empirical MEG data. Hyperparameters for spectral modelling were as follows: a minimum peak height of 0.1 a.u., a maximum of 6 peaks, peak width limits between [1, 12] Hz, and a Gaussian overlap threshold of 0.75 s.d. for the empirical dataset and 2.0 for the simulated data. The choice of hyperparameter settings was informed by visual inspection of the data. The most parsimonious spectral model was selected according to the Bayesian Information Criterion.

Note that all three methods relied on the same *specparam* modelling fitting procedure, resulting in the same model fits for all three simulations, and equivalent errors in arrhythmic parameter estimates. Differences in the recovered relationships between arrhythmic exponent and rhythmic alpha power across the three methodologies, therefore, cannot be explained by differences in the *specparam* model fit.

### Empirical dataset

Data of 606 participants from the Cambridge Centre for Aging and Neuroscience repository (CamCAN) were used to validate our results obtained in simulation (mean age = 54.69, SD = 18.28; 299 female) (Taylor et al., 2017). Each participant completed a resting-state, eye-closed MEG recording using a 306-channel VectorView MEG system (MEGIN, Helsinki, Finland) that lasted approximately 8 minutes. MEG data were collected using 102 magnetometers and 204 planar gradiometers sampled at 1 kHz with a 0.03-330 Hz bandpass filter.

### MEG preprocessing

We preprocessed the magnetoencephalography (MEG) recordings using Brainstorm (March 2021 distribution) (Tadel et al., 2011) in MATLAB (2020b; Natick, MA), adhering to established best-practice guidelines (Gross et al., 2013). The preprocessing methodology followed the protocols detailed previously in (Castanheira et al., 2024; da Silva Castanheira et al., 2021; L. E. Wilson et al., 2024).

Data were filtered to remove i) line noise artifacts at 50 Hz and its first 10 harmonics, ii) an 88-Hz artifact present in the Cam-CAN dataset (Wiesman, da Silva Castanheira, et al., 2022), and iii) slow-wave and DC-offset artifacts using a high-pass finite impulse response (FIR) filter with a cutoff frequency of 0.3 Hz. To mitigate the impact of cardiac artifacts, low-frequency (1–7 Hz) and high-frequency (40–400 Hz) artifacts, we applied Signal-Space Projections (SSPs) and removed the first projector, which explained the most variance.

Neural time series were co-registered to the individual T1-weighted MRI of each participant, facilitated using approximately 100 digitized head points. Biophysical head models were computed using *Brainstorm*’s overlapping-spheres model (default parameters), and source models were computed using linearly constrained minimum-variance (LCMV) beamforming following *Brainstorm*’s default parameters (2018 version for source estimation processes). Orientations of the 15,000 sources were constrained normal to the cortical surface. Neural power spectra were then computed using the Welch’s method, utilizing 2-second windows with a 50% overlap for every region of the Destrieux atlas (Destrieux et al., 2010).

### Statistics

We calculated Spearman correlations and 95% confidence intervals using the DescTools package in R (R Core Team, 2022). We computed the Spearman correlation between rhythmic power and arrhythmic exponent in Figure 2, between rhythmic alpha and beta power in Figure 4, between rhythmic alpha power and exponent in Figure 5, and between simulated demographics and alpha power in Figure S4.

We computed the error in arrhythmic parameter estimates by subtracting the ground truth parameters used in simulating the neural time series from their estimated values. We then assessed the relationship between the error in arrhythmic parameters and estimated alpha power using regression models implemented in R. The results of these analyses are presented in Figure 4.

## Results

We simulated 16,000 neural time series across five experiments with known ground truth relationships between different spectral parameters of interest. We then asked how well three different methods for quantifying rhythmic brain activity could recover these simulated relationships. These three methods were denoted i) modelled rhythmic power, ii) linear-detrended power and iii) log-detrended power averaged within a narrow band (see Methods for details). For simplicity, we will refer to the detrending methods averaged within a narrow band as detrended power. The key difference between the three methods is that modelled rhythmic power explicitly uses the modelled Gaussian height of peaks that sit above the 1/f arrhythmic background activity from the outputs of the *specparam*. In contrast, the detrending power methods remove the estimated 1/f arrhythmic power from the spectrum, either in log or linear space. The detrending methods do not rely on the modelled Gaussian peaks, but instead estimate rhythmic power from the residual variance after removing the arrhythmic background activity from the power spectrum. Note that all three methods detrend the power spectrum, yet the key difference between the methods is in how rhythmic power is computed.

### Spectral detrending introduces spurious correlations between arrhythmic and rhythmic components

In our first set of simulations, we evaluated the accuracy of each method in independently recovering the arrhythmic exponent and rhythmic alpha power. In these simulations, there was no ground truth correlation between these components (i.e., ⍴ = 0). As *specparam* characterises the contribution of Gaussian peaks independently of the 1/f background noise, we expected that modelled power would best recover this null relationship between the two signal components. In contrast, as detrending approaches depend on first modelling the neural power spectrum to accurately remove the contribution of the 1/f background noise, we hypothesized that this may introduce spurious correlations between the two signal components due to errors in model fit. We ran three sets of simulations varying i) alpha centre frequency, ii) alpha amplitude, and iii) arrhythmic exponent.

For the alpha centre frequency simulations, we observed that the choice of method to quantify rhythmic activity substantially impacted the recovered relationship between rhythmic alpha power and arrhythmic exponent. For modelled power, the observed relationship between alpha power and exponent was generally close to the ground truth correlation of 0. Spearman correlation values were less than |0.15| for every simulated centre frequency except for 7 and 8 Hz, where we observed weak spurious relationships between alpha power and exponents (⍴ = -0.18, CI [-0.27, -0.09]; -0.15 (CI [-0.24, -0.06], respectively) (Figure 2a left panel). Note that this difference in estimated correlation was not driven by a worse model fit (Figure S1 & S2). In comparison, we observed systematic spurious negative correlations between rhythmic and arrhythmic components (Spearman correlation values below -0.15) for both log- and linear-detrended methods. Log-detrended power performed the poorest of the three methods, with an average negative correlation between rhythmic alpha power and arrhythmic exponent of ⍴ =-0.35 (Figure 2a, middle panel). In contrast, the average estimated relationship between rhythmic alpha power and the arrhythmic exponent was ⍴ = -0.17 for linear-detrended power and ⍴ = - 0.03 for modelled power (Figure 2a, left and right panels).

Next, we evaluated how well the three methods were able to recover a ground truth null relationship of alpha power and arrhythmic exponent for various simulated values of arrhythmic exponents, holding alpha centre frequency and amplitude constant (see Methods for details). Of the three methodologies for quantifying rhythmic alpha power, modelled power was the closest to ground truth with an average correlation between rhythmic alpha power and arrhythmic of ⍴ = 0.03, in comparison to ⍴ =-0.12 for linear-detrended power and ⍴ =-0.39 for log-detrended power (Figure 2b). The value of the simulated arrhythmic exponent did not substantially impact the estimated relationship between exponent and alpha power for log-detrended power. These effects are not specific to the alpha band, but are similarly observed when exploring the relationship between rhythmic beta power and the arrhythmic exponent (see Figure S3). A similar pattern of parameter recovery was obtained when varying alpha amplitude, reported in Supplemental Materials (see Figure S4). Taken together, these analyses suggest that modelled power is the most robust method for quantifying rhythmic alpha power independently of arrhythmic brain activity.

Given the spurious relationships obtained between rhythmic alpha power and arrhythmic exponent, we next explored whether methodological choices in quantifying rhythmic brain activity may introduce similarly spurious relationships between different brain rhythms. To this end, we explored recovery of linear monotonic relationships between estimates of rhythmic alpha (8-12 Hz) and beta power (12-24 Hz) for the three methods of computing rhythmic power. Akin to the previous results, we observed spurious positive correlations between alpha and beta amplitudes for log-detrended power (*log-log* mean ⍴ = 0.54) and linear-detrended power (*linear* mean ⍴ = 0.44; see Figure 2c). In contrast, for modelled power, alpha and beta power were generally unrelated to one another across simulations of various arrhythmic exponents (mean ⍴ = 0.11) and therefore closer to the ground truth of zero correlation.

The spurious relationship between rhythmic alpha and beta power increased in strength for linear-detrended power (Figure 2c, right panel); we hypothesised that this was due to an increasing influence of background signals on neural activity with larger arrhythmic exponents. In other words, at larger arrhythmic exponents, the total amount of variance in the power spectrum driven by the arrhythmic exponent in linear space appears to be proportionally larger than its rhythmic components. To substantiate these claims, we ran a mediation analysis to test whether the estimated arrhythmic exponent (M) mediated the relationship between linear-detrended beta power (X) and alpha power (Y). Consistent with our hypothesis, the bootstrapped average causal mediation effect was significant (0.26, 95% CI [0.23, 0.29], p < 0.001), indicating a partial mediation explaining 31% of the total effect.

One issue in interpreting our simulation results is that the detrending methods rely on averaging power within a narrow frequency band, while the modelled rhythmic power relies on a single point parameter estimate. To address this concern, we defined alpha power as the highest values in a narrow band (8-12 Hz) after detrending and computed the relationship between alpha power and arrhythmic exponent for our simulations, where we should observe no systematic relationship between the two. We observed similar spurious relationships between alpha power and arrhythmic exponent using these point estimates of detrended alpha power (see Figure S5).

We ran another set of simulations where we systematically manipulated the value of alpha peak bandwidth, holding all other spectral parameters constant (Figure 3a). Of the three methodologies for quantifying rhythmic alpha power, linear detrended power was the closest to the ground truth, with an average correlation between rhythmic alpha power and arrhythmic of ⍴ = -0.20, in comparison to ⍴ = -0.28 for modelled rhythmic power, and ⍴ = -0.57 for log-detrended power (Figure 3b). We note that modelled rhythmic power independently recovered arrhythmic exponent and rhythmic power for alpha peak bandwidths below 3 Hz (Figure 3b). We similarly observed that error in parameter estimates for the arrhythmic exponent was significantly more variable for alpha peak bandwidths above 3Hz (Figure 3c). These findings suggest that for large broad spectral peaks, *specparam* may struggle to accurately recover the arrhythmic exponent at the simulated signal-to-noise level, leading to poorly estimated arrhythmic slopes and resulting in spurious correlations between the arrhythmic and rhythmic model parameters.

### Error in spectral parameter estimates introduces spurious correlations between signal components

Next, we explored why detrending approaches—particularly in log space—result in spurious correlations between spectral parameter outputs. We chose to focus our analyses on log-detrended power as it is the method with which we see the largest spurious relationships between arrhythmic exponents and rhythmic alpha power. We hypothesized that errors in the estimated arrhythmic model parameters could explain the observed relationships between rhythmic and arrhythmic model parameters. To test this hypothesis, we fit a linear regression model where we predicted estimates of detrended power from the error in the arrhythmic parameter estimate (see Methods for details).

We observed that for spectra in which the arrhythmic offset was underestimated, log-detrended alpha power was also estimated as larger (β =0.30, SE =1.36 * 10^-2^, *p* < 0.001, CI [ 0.27, 0.32]), with the largest errors in estimates of offset yielding alpha power estimates closest to the ground truth (dashed line Figure 4a top panel). We attribute this effect to the underestimation of alpha power using log-detrending approaches (i.e., most points fall below the dashed horizontal line in Figure 4a). In contrast, error in estimates of the arrhythmic slope showed a nonlinear quadratic relationship to log-detrended alpha power, for which the larger the absolute error in arrhythmic exponent, the higher the alpha power estimate (β =0.63, SE =1.09 * 10^-2^, *p* < 0.001, CI [ 0.61, 0.65]). Error in both arrhythmic parameters independently predicted alpha power estimates (offset: β = 0.08, SE =1.25 * 10^-2^, *p* < 0.001, CI [ 0.06, 0.11]; exponent: β = 0.61, SE =1.15 * 10^-2^, *p* < 0.001, CI [ 0.59, 0.63]).

We observed weak significant relationships between modelled alpha power and error in arrhythmic parameter estimates (offset: β = -0.08, SE =1.68 * 10^-2^, *p* < 0.001, CI [ -0.13, -0.06]; exponent: β = 0.15, SE =1.55 * 10^-2^, *p* < 0.001, CI [ 0.12, 0.18]; see Figure 4a bottom panel). Note that although these relationships were statistically significant, they explained only a modest amount of total variance in modelled alpha power (3% of the variance in comparison to 42% of the variance in detrended alpha power), explaining why this method was largely successful in independently recovering these components.

Two example power spectra can be used to illustrate this effect. In the top panel of Figure 4b, a simulated neural power spectrum has been correctly modelled with two rhythmic peaks in the alpha and beta range, as well as an appropriate arrhythmic model. As a result, the detrended spectrum, plotted as the inlaid line graph, is unbiased. In contrast, when arrhythmic parameter estimates are incorrectly estimated as depicted in the bottom panel of Figure 4b, the detrended rhythmic power is greater than expected (Figure 4b bottom panel inlaid graph) – in this example, rhythmic power in lower frequencies is overestimated because of the underestimation of the arrhythmic offset. More specifically, the modelled arrhythmic offset is estimated to be lower than the ground-truth simulated arrhythmic offset, resulting in larger estimates of rhythmic power for detrending methods. Detrending analyses are significantly more sensitive to errors in the arrhythmic model and, as a result, may introduce spurious relationships between estimated rhythmic and arrhythmic components.

### Handling instances where *specparam* fails to model a peak

A specific disadvantage of modelled power, relative to detrending methods, is that it introduces missing values when the *specparam* algorithm fails to fit a Gaussian peak within a pre-defined narrow band frequency range (e.g., the alpha band). We therefore sought to explore the influence of these missing values on the recovered relationship between arrhythmic exponent and alpha power. We simulated 100 neural time series for 10 effect sizes describing the relationship between alpha power and arrhythmic exponent (from ⍴ = 0.00 to ⍴ = 0.90 in steps of 0.10). We observed that modelled power, on average, recovered the ground truth simulated relationship between alpha power and arrhythmic exponent (Figure 5a). We note, however, that larger effect sizes (⍴ > 0.50) were systematically underestimated. We then tested the influence of replacing missing peak amplitudes not modelled by *specparam* with a value of 0.00. On average, estimated relationships between alpha amplitude and arrhythmic exponent were closer to the ground truth when missing peak amplitudes were replaced with zeros (Figure 5b).

Log-detrended power was similarly capable of recapitulating the simulated relationship between alpha power and arrhythmic exponent (Figure 5c). In contrast, linear-detrended power systematically overestimated the relationship between arrhythmic exponent and alpha power, principally for weaker relationships (Figure 5d). The mean absolute error between the simulated relationship and estimated correlation was 0.26 for modelled rhythmic power, 0.21 for log detrended power, and 0.31 for linear detrended power. The spurious relationships between the arrhythmic exponent and rhythmic alpha power observed for log detrending methods (see Figure 2) resulted in estimated correlations that were, on average, higher than modelled power. In contrast, modelled power tended to underestimate the simulated relationships between arrhythmic slope and rhythmic alpha power (see Figure 5a&b)—explaining why log detrending methods achieved slightly lower errors between the simulated and estimated relationships.

Note that all three methods relied on the same *specparam* modelling fitting procedure, resulting in the same model fits for all three simulations, and equivalent errors in arrhythmic parameter estimates. Differences in the recovered relationships between arrhythmic exponent and rhythmic alpha power across the three methodologies, therefore, cannot be explained by the *specparam* model fit.

### Modelled vs detrended rhythmic power leads to diverging interpretations

Taken together, our simulation analyses corroborate our hypothesis: modelled power outperforms detrending methods and accurately quantifies the independent contributions of rhythmic and arrhythmic brain activity. We next sought to verify our results obtained in simulation in an empirical dataset. In particular, our goal was to highlight how different methods of quantifying rhythmic amplitude may yield diverging interpretations. We analyzed the relationship between alpha amplitude and arrhythmic exponent, two proposed markers of cortical inhibition, in the resting-state MEG data of 606 individuals within the CamCAN database (18-89 years old).

We estimated the linear monotonic relationship between resting-state arrhythmic exponents and alpha power using Spearman correlations for each of 148 cortical parcels of the Destrieux atlas. We observed a rostro-caudal gradient, with frontal brain regions showing a positive linear relationship and posterior areas having a negative relationship between log-detrended alpha power and arrhythmic exponent (Figure 6a, left panel). In contrast, the relationship between modelled alpha power and arrhythmic exponent was positive across the entire cortex, except within nine occipital parcels (Figure 6a left panel). See Figure S6 for FDR thresholded brain maps of this relationship. We then computed the linear relationship between alpha power and arrhythmic exponent for every parcel across three age groups (young adults 18-45, adults 45-65, and older adults 65+ years old). Notably, we observed that the method used to quantify rhythmic alpha amplitude meaningfully impacted the results (Figure 6b). Using the log-detrended power approach, we found that the relationship between alpha power and the arrhythmic exponent varied depending on the age group assessed. Older adults showed more positive correlations between alpha power and arrhythmic exponent compared to younger adults – an effect which was most pronounced in fronto-central and temporal brain regions. However, this dependency of spectral relationship on age disappeared for modelled alpha power, suggesting it may be a spurious consequence of the detrending approach (Figure 6b, bottom row).

**Figure 6:**
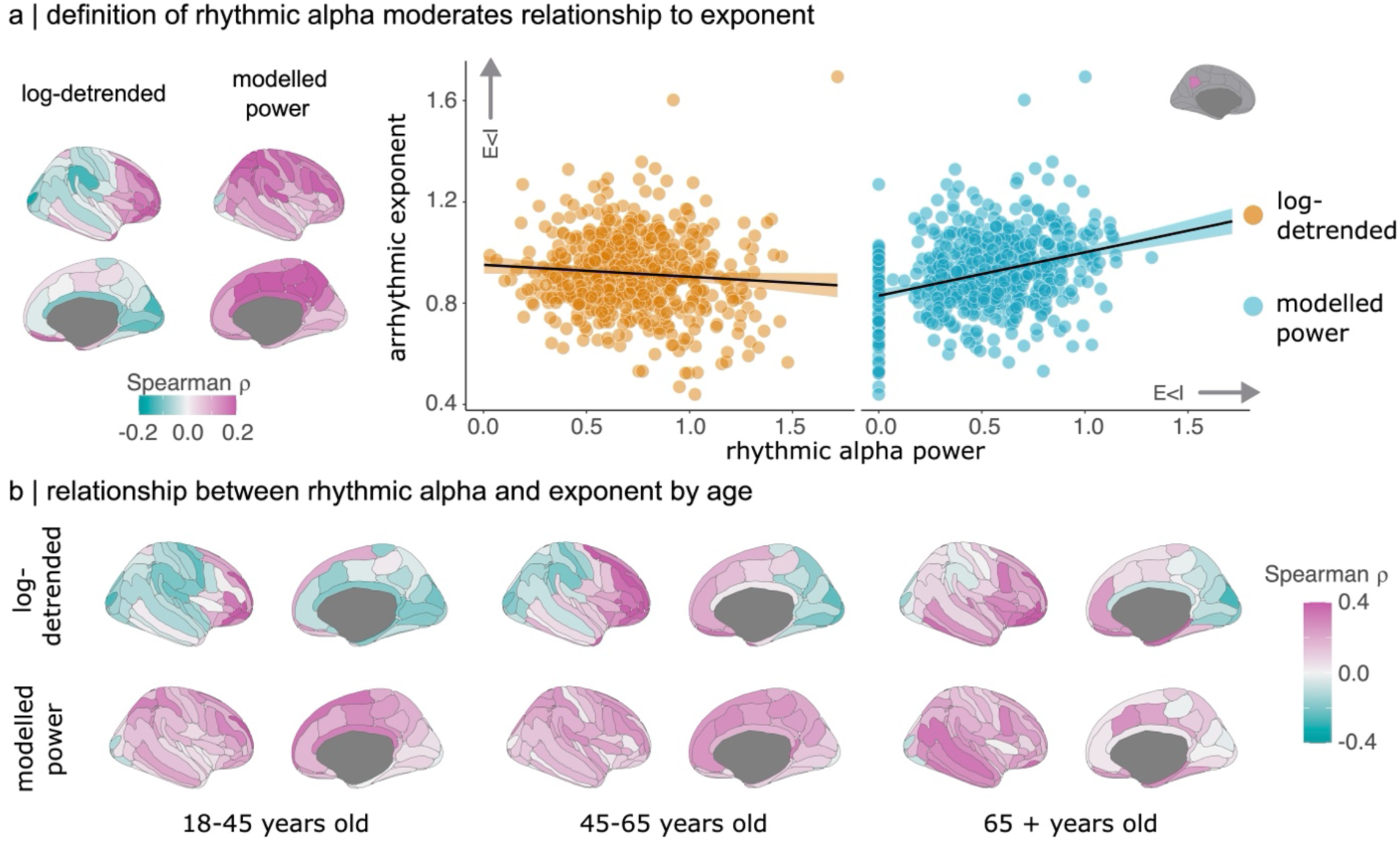
The choice of method for quantifying rhythmic power impacts obtained relationships between spectral components (empirical data). (a) Left panel: Topographic map of the linear monotonic relationship between rhythmic alpha amplitude and arrhythmic exponent for log-detrended (left) and modelled alpha power. Log-detrended alpha power negatively relates to arrhythmic exponent, principally in posterior brain regions, including the visual cortex. In contrast, modelled alpha power positively relates to arrhythmic exponent across the entirety of the cortex, except for nine occipital regions. Right panel: Scatter plot of the relationship between rhythmic alpha power and arrhythmic exponent for an exemplar brain region. While log-detrended alpha power weakly negatively predicts arrhythmic exponent, modelled alpha power positively predicts arrhythmic exponent in this brain area. Note that the distribution of zero alpha amplitude values reflects participants for whom *specparam* did not model a Gaussian peak. (b) Topographic map of the linear monotonic relationship between rhythmic alpha amplitude and arrhythmic exponent for i) different age groups (columns) and ii) the two methods for quantifying rhythmic power (rows). The interpretation of the relationship between resting-state alpha power and arrhythmic exponent differs substantially depending on the method for quantifying the amplitude of alpha power.

We also examined the impact of these methodological choices on other individual differences. First, we observed strong positive relationships between rhythmic alpha and beta amplitudes when using log-detrended power – relationships which were not obtained for modelled power (Figure S7). When evaluated in combination with our simulation results, these findings suggest that detrending approaches may introduce spurious correlations between brain rhythms. Second, we computed the relationship between alpha amplitude and chronological age, and observed that it, again, depended on the method of quantifying power (see Figure S8). Third, we observed that the relationship between the arrhythmic exponent and frontal log-detrended beta power inverts when relying on modelled beta power (see Figure S9). Together, our findings in the CamCAN dataset illustrate that the choice of method for quantifying rhythmic activity can yield diverging and potentially spurious results when applied to empirical data.

## Discussion

The analysis of brain rhythms is central to the field of neuroscience. For over a century, researchers have sought to establish links between neural oscillations, behaviour, and disease (Mushtaq et al., 2024). However, methods for quantifying rhythmic power vary considerably and make differing assumptions (Brake et al., 2024; Donoghue et al., 2020; Wen & Liu, 2016). We explored how the choice of method for computing the amplitude of brain rhythms impacts the recovery of simulated ground-truth relationships between different spectral features. Addressing this question is critical to establishing whether rhythmic and arrhythmic signal components differentially relate to behaviour. We demonstrate, based on simulated neural time series, that modelling Gaussian amplitudes (e.g. as implemented in *specparam*) provides a robust method for quantifying the power of brain rhythms independent of the arrhythmic exponent. Modelled power recovers independent sources of variance that are minimally contaminated by arrhythmic signal components. In contrast, linear and log detrending methods can introduce spurious relationships between arrhythmic and rhythmic parameters.

### Is spectral detrending necessary?

Previous research has treated background 1/f brain activity as noise to be removed when quantifying the amplitude of brain rhythms. Spectral detrending or whitening is used to correct for this background activity, either by computing relative power or modelling the arrhythmic signal component (Donoghue et al., 2020). Despite the growing use of algorithms to parameterize the neural power spectra (Donoghue et al., 2020; L. E. Wilson et al., 2022, 2024) and estimate arrhythmic signal components (Wen & Liu, 2016), questions remain about whether researchers should detrend the power spectrum when clear oscillations are present. Indeed, recent computational modelling work suggests that spectra require different types of detrending—i.e., dividing vs subtracting—depending on the specific physiological mechanisms one assumes to drive observed changes in the arrhythmic spectrum (Brake et al., 2024). Brake et al. (2024) concluded that spectral detrending should be avoided altogether unless a clear biophysical and physiological justification is provided. Yet the rhythmic components of the neural power spectra were not explicitly modelled using Gaussian peaks, as pursued in *specparam*. By ignoring dynamics in background brain activity, researchers may conflate arrhythmic signal components with narrow-band modulations of power. This latter point is of particular interest when testing for potential physiological correlates of behaviour.

Background 1/f activity has historically been treated as noise which needs to be corrected (Donoghue et al., 2020, 2022; Ouyang et al., 2020; Wen & Liu, 2016). The reasoning was that 1/f arrhythmic activity could obscure statistical differences in narrowband power estimates and introduce false-positive narrowband changes. To address some of these concerns, previous model-free approaches converted neural power to decibel (dB) units by dividing power spectra by a baseline and log-transforming the resulting spectrum. Recent findings have questioned this practice (Gyurkovics et al., 2021), suggesting that the decibel conversion requires untenable assumptions which can distort statistical findings when they are not met.

The present paper explores this question with ground-truth simulations. Our findings suggest that modelling rhythmic activity by fitting Gaussian peaks recovers the simulated amplitude of neural oscillations with greater accuracy than detrending approaches, which are subject to biases (Figure 2). These biases are introduced when the arrhythmic component of the power spectrum is not accurately fit, leading to spurious correlations between model parameters (Figure 4). The spurious relationships observed in detrending analyses are not limited to rhythmic and arrhythmic exponents. We found that the amplitudes of different rhythmic oscillations are spuriously correlated if the background 1/f component of the spectrum is not adequately detrended (Figure 2c). This poses challenges for isolating the independent contribution of multiple brain rhythms.

### The relationship between arrhythmic and rhythmic brain activity

Alpha power is theorized to reflect cortical inhibition, principally in the sensory cortex (Clayton et al., 2018; Foxe & Snyder, 2011; Jensen, 2024; Jensen & Mazaheri, 2010; Morrow et al., 2023; Samaha et al., 2020b). Phases of alpha power are believed to reflect periods of local inhibition and disinhibition, with increases in alpha amplitude believed to reflect increased cortical inhibition. Posterior alpha rhythms similarly predict the likelihood of reporting phosphenes triggered by transcranial magnetic stimulation (TMS), which is interpreted in support of the inhibition interpretation of alpha rhythms (Romei, Brodbeck, et al., 2008; Romei, Rihs, et al., 2008). Work on the orienting of attention corroborates this theory, with decreased alpha power observed contralateral to the attended visual hemifield (Bagherzadeh et al., 2020; Jensen, 2024; Landry et al., 2024).

A separate line of work has proposed that the arrhythmic exponent reflects the relative contribution of excitatory and inhibitory populations of neurons. Several lines of evidence support this view: computational work suggests that the arrhythmic exponent reflects the relative contribution of AMPA and GABA_A_ populations (Gao et al., 2017). Similarly, administration of anesthetics alters the slope of the arrhythmic component (Colombo et al., 2019; Lendner et al., 2020; Maschke et al., 2023; Waschke et al., 2021). Recent work leveraging a pupil-based biofeedback paradigm demonstrates that changes in the brain’s state of arousal are correlated to shifts in cortical arrhythmic brain activity (Weijs et al., 2025).

Our analysis of empirical data demonstrates how the interpretation of the relationship between alpha power and the arrhythmic exponent depends on the methodological choices. Log-detrending methods introduce a spurious negative relationship between arrhythmic exponents and alpha power signals at rest (see Figure 6a). In contrast, when using spectral modelling, we observed that alpha power and arrhythmic exponent are positively correlated (Figure 6b) across the adult lifespan, in line with what we would expect from theories of cortical inhibition. This observation dovetails with recent findings suggesting that the two signal components are positively correlated during a perceptual decision-making task (Elliott et al., 2025).

While our results speak to the shared variance between two hypothesized markers of inhibition, a substantial portion of variance remains to be explained. We speculate that different biophysical parameters may govern local inhibition (i.e., the conductance of channels versus the activity of inhibitory cell populations), which would explain why we did not observe that alpha power and the arrhythmic exponent are not more strongly correlated with one another. In line with this interpretation, Brake and colleagues demonstrated using computational modelling that arrhythmic exponents may not be a reliable marker of E-I balance, given the nonlinearities of membrane dynamics (Brake et al., 2024). It is therefore likely that both signals reflect a cascade of complex neuronal functions, some of which reflect cortical inhibition. Future empirical and computational work should seek to clarify the biophysical underpinnings of both alpha rhythms and background 1/f activity.

This literature suggests that the arrhythmic exponent and alpha power should be closely related, given the close link between cortical inhibition and the E-I balance. However, empirical findings on the relationship between narrowband alpha power and arrhythmic exponent have proven inconsistent. Gyurkovics and colleagues found little evidence for a relationship between resting-state alpha power and the 1/f slope (Gyurkovics et al., 2021). In contrast, Muthukumaraswamy and Liley found strong positive relationships across the cortex (Muthukumaraswamy & Liley, 2018). These inconsistent findings may be driven by the variation in methodological choices we document here (see Figure 6). Further work is needed to clarify whether narrowband changes in power are related to broadband changes in the arrhythmic exponents, as would be predicted by a multiplicative model of neural power spectra.

Despite the theoretical interpretation of alpha power and the arrhythmic exponent as markers of cortical inhibition, *specparam* and related models assume that the contributions of arrhythmic and rhythmic signal components are independent (additive) of one another (Donoghue et al., 2020; Medrano et al., 2025; L. E. Wilson et al., 2022, 2024). Accordingly, our findings indicate that modelling Gaussian amplitudes can recover the independent contribution of arrhythmic and rhythmic signal components in simulation (Figure 2). However, future work may fruitfully explore whether a comprehensive neurophysiological generative model could explain the empirical relationships observed between alpha power and arrhythmic exponent at rest.

### Age-related shifts in arrhythmic and rhythmic brain activity

Both rhythmic and arrhythmic neurophysiological activity shifts across the lifespan. For example, the alpha rhythm slows in older age, reflecting functional decline (Babiloni et al., 2006; Merkin et al., 2022; Moretti et al., 2013; Scally et al., 2018; Thuwal et al., 2021). Similarly, the aperiodic signal component evolves across the lifespan and is associated with age-related declines in sensory and cognitive functions (He et al., 2019; Karalunas et al., 2022; Schaworonkow & Voytek, 2021; Thuwal et al., 2021; Voytek et al., 2015b; Voytek & Knight, 2015). These age-related changes in both arrhythmic and rhythmic brain activity are a part of normal aging, but have also been observed in neurological diseases (Gallego-Rudolf et al., 2024; Wiesman et al., 2023, 2024). Further work is needed to differentiate between normal age-related changes in neurophysiology and pathological changes in brain activity.

The release of large open datasets has made the possibility of lifespan investigations of rhythmic and arrhythmic brain activity increasingly attainable (Da Silva Castanheira et al., 2025; Taylor et al., 2017). Akin to efforts harnessing MRI (Bethlehem et al., 2022), growth charts of arrhythmic and rhythmic activity may represent an important step in monitoring brain health across the lifespan and developing personalized medicine approaches. To achieve such goals, it is important to understand whether the observed rhythmic and arrhythmic changes in brain activity are independent of one another or share a common neural mechanism. Our findings support a hypothesis that alpha rhythms and arrhythmic exponents are positively related at rest across the lifespan. We speculate that this coupling reflects overlapping neural mechanisms underlying age-related changes in arrhythmic and rhythmic signals. To substantiate such claims, longitudinal research is needed to disentangle how arrhythmic and rhythmic signals change across the lifespan.

### The independent contribution of rhythmic and arrhythmic components to behaviour

There is growing evidence that both arrhythmic and rhythmic signal components contribute to behaviour in health and disease (Koenig & He, 2025; Wiesman et al., 2023). However, establishing the relative contribution of these components remains a nascent project, one in which requires the development of robust methods such as the ones we outline here.

Take for example, the field of metacognition research, where evidence suggests that both arrhythmic and rhythmic power, measured prior to stimulus onset, can predict participants’ subjective visibility and confidence reports over and above variation in objective performance (Benwell et al., 2017; Cunningham et al., 2023; Limbach & Corballis, 2016; Samaha et al., 2017, 2020b, 2022; Wöstmann et al., 2019). In light of our present analyses, it will be important to test for potentially complex relationships between arrhythmic signals and alpha power, and explore a hypothesis that each component may make a distinct contribution to metacognition.

We argue that understanding how these two signal components contribute to behaviour, and whether their contributions are independent of one another, is an important next step in advancing our understanding of the underlying neural mechanisms. In our paper, we document how methodological choices may impact the independent estimation of arrhythmic and rhythmic brain activity. This will have important implications for researchers who aim to explore such questions.

### Methodological Considerations

Based on our simulations, we recommend that researchers explicitly model Gaussian amplitudes when quantifying rhythmic power, using algorithms such as *specparam*. We show that this method minimizes the potential for spurious relationships between arrhythmic and rhythmic model parameters (Figure 2). One methodological consideration when using modelled power is the large number of missing data points that occur as a result of *specparam* failing to fit a spectral peak within a predefined narrow-band frequency range (e.g., alpha 8- 12 Hz). We investigated whether this disadvantage can be overcome by replacing the missing values with an amplitude of zero (e.g., Figure 5). Our simulations suggest that this approach accurately recovers simulated relationships. We note, however, that this analytic choice may alter the distribution of the resulting data, rendering non-parametric statistics necessary. In addition, this approach of dealing with missing data only applies to the amplitude of rhythmic oscillations. In the case of center frequencies, for example, it remains unclear what the best approach is to deal with missing data.

Another methodological consideration is the use of pre-defined fixed ranges to delineate frequency bands of interest (e.g., alpha band 8- 12 Hz). There is considerable inter-individual diversity in brain rhythms, including the specific centre frequency of the rhythm. This makes aggregating data across individuals challenging, specifically if an individual’s centre frequency falls at the edge of the canonical frequency band definitions. We observed in our simulations that centre frequencies at the lower edge of the pre-defined narrow-band may introduce spurious relationships between narrow-band estimates of rhythmic power and the arrhythmic exponent (see Figure 2a, modelled power). While these spurious relationships are much smaller in magnitude than the ones observed with detrending methods, we would advise researchers to explore inter-individual differences in the centre frequency of brain rhythms before defining a frequency band of interest.

We simulated spectral peaks of varying bandwidths and observed spurious correlations between the arrhythmic exponent and estimates of rhythmic power for all three methodologies, principally for broad spectral peaks (i.e., bandwidths > 3 Hz). We observed that modelled rhythmic power recovered ground truth simulations for narrow spectral peaks (see Figure 3). We interpret these findings as follows: broad spectral peaks challenge the *specparam* fitting procedure at the simulated SNR levels. This can result in large errors in the estimates of the arrhythmic slope, which in turn introduce spurious relationships between the spectral parameters. We therefore recommend visually inspecting the quality of the fitted spectra before hypothesis testing, particularly if the spectra contain broad peaks, to ensure the accuracy of *specparam* model fits.

In the case where a hypothesis concerns a specific frequency (e.g., a 10 Hz oscillation), we similarly observed that spectral detrending methods can result in spurious relationships (see Figure S5). This suggests that the issues introduced by spectral detrending methods are not related to averaging the residual variance across a spectral band, but instead directly related to how rhythmic power is estimated from the residual variance.

In our simulations, we did not vary the frequency range over which to detrend the data, which may affect our results. Instead, in line with previous recommendations (Donoghue et al., 2020, 2022), we suggest modeling the neural power spectra using *specparam* over the largest frequency range in which the background arrhythmic activity remains linear in log-log space (i.e., most similar to a 1/f decay). This adheres to *specparam’s* assumptions and ensures that the model is fit over as much data as possible. Proper estimation of the 1/f arrhythmic component is critical for the independent estimation of rhythmic and arrhythmic components (see Figure 4). As we observed in our simulations (see Figure 4), errors in the estimates of the 1/f background activity may lead to spurious correlations between the arrhythmic exponent and rhythmic power. Minimizing fitting error of the *specparam* model, therefore, is critical to obtaining independent estimates of arrhythmic and rhythmic spectral parameters.

Spectra across large frequency ranges often contain spectral bends, referred to as ‘knees’ in the literature (Donoghue et al., 2020). Our current analyses did not explicitly test whether spectral bends impact the quantification of rhythmic brain activity. This choice was informed by practical constraints, given our aim was to build on and evaluate existing algorithms such as *ms-specparam*, which does not fit spectral bends (L. E. Wilson et al., 2024). It is likely that improper modelling of spectral bends in the arrhythmic component will similarly lead to spurious relationships between arrhythmic model parameters and estimates of rhythmic power that we document here, and this potential confound could usefully be explored in future work.

Our simulated data were generated with the NeuroDSP toolbox (Cole et al., 2019), which sums the time series of the simulated rhythmic and arrhythmic components and generates synthetic time series data without bursting dynamics and other non-linearities. This choice may in fact favour log-detrending methods, as it relies on log-summed power to generate the time series. It is therefore notable that we find that log-detrended power exhibited the largest spurious correlations between arrhythmic and rhythmic components in our simulated datasets. Future work should investigate how non-linear properties of the time series, such as bursting, impact the estimation of arrhythmic and rhythmic brain activity.

Our paper focused on the *specparam* family of spectral detrending methods, this choice was informed by the widespread use of *specparam* and its related methods (Donoghue, 2024; Donoghue et al., 2020; L. E. Wilson et al., 2022, 2024) and the explicit parametric modelling of spectral peaks. We did not explore how other spectral detrending methods like IRASA, the α +1/f method, or simply fitting a linear model may impact the results. We speculate that any spectral detrending method which does not explicitly model spectral peaks will show similar results to the reported detrending methods in our present paper. We come to this conclusion as the primary source of spurious correlations between brain rhythms and the arrhythmic exponent was the inaccurate modelling of the 1/f background activity. Under the assumption that methods to separate the 1/f from neural oscillations all work equally well, the choice of method for quantifying brain rhythms should be negligible. This, however, is not the case. For example, previous work reports worse arrhythmic exponent fits with IRASA in comparison to *specparam* in the presence of broad spectral peaks (McKeown et al., 2024). Taken together, there is considerable interest in the field to separate the independent contributions of arrhythmic and rhythmic signal components, yet the methods to do so are still in their infancy. Future work should continue to explore how methodological choices impact the interpretation of results.

In conclusion, we demonstrate that analytical choices when quantifying the amplitude of brain rhythms can lead to spurious correlations between arrhythmic and rhythmic spectral parameters and confound interpretations of findings. By relying on simulations of neural time series, we demonstrated that detrending approaches lead to misleading negative correlations between the arrhythmic exponent and alpha power. In contrast, modelled power provides the most robust approach for quantifying the amplitude of brain rhythms. Based on modelled Gaussian peaks, we show that arrhythmic exponents and alpha power are positively correlated across the cortex in individuals sampled from a large age range. In line with theories suggesting that both resting state alpha power and the arrhythmic slope reflect cortical inhibition, our results suggest that both signal components share a considerable amount of variance. These findings, in turn, raise new questions about whether neural spectra are best captured by a multiplicative or additive model. We anticipate that the findings of this paper will inform future research on the neural mechanisms of these processes, encouraging more robust, reproducible, and interpretable results.

## Data & code availability

All in-house code used for data analysis and visualization is available on GitHub https://github.com/jasondsc/IndependenceArrhythmicRhythm. The reanalyzed data presented herein are available from https://cam-can.mrc-cbu.cam.ac.uk. The simulated neural time series are available upon reasonable request from the first author.

## Author Contributions

Conceptualization: J.d.S.C., M.L., and S.F.

Data Curation: J.d.S.C.

Methodology: J.d.S.C., M.L., and S.F.

Software: J.d.S.C.

Visualization: J.d.S.C., and S.F.

Validation: J.d.S.C.

Formal analysis: J.d.S.C.

Supervision: M.L., S.F.

Project administration: S.F.

Writing—original draft: J.d.S.C. and S.F.

Writing—review and editing: J.d.S.C., M.L., and S.B.

## Supplemental Materials

**Table S1.** Error in estimating arrhythmic spectral parameters predicts log-detrended alpha power.

| *Predictors* | Log-detrended alpha power |  |  |
| --- | --- | --- | --- |
|  | *Estimates* | *CI* | *p* |
| (Intercept) | 0.00 | -0.02 – 0.02 | 1.000 |
| Error in offset parameter | 0.08 | 0.06 – 0.11 | **<0.001** |
| Error in exponent parameter (1<sup>st</sup> order) | 0.12 | 0.10 – 0.14 | **<0.001** |
| Error in exponent parameter (2<sup>nd</sup> order) | 0.61 | 0.59 – 0.63 | **<0.001** |
| Observations | 4935 |  |  |
| R<sup>2</sup> / R<sup>2</sup> adjusted | 0.418 / 0.417 |  |  |

**Table S2.** Error in estimating arrhythmic spectral parameters weakly predicts modelled alpha power.

| *Predictors* | Modelled alpha power |  |  |
| --- | --- | --- | --- |
|  | *Estimates* | *CI* | *p* |
| (Intercept) | -0.00 | -0.03 – 0.02 | 0.796 |
| Error in offset parameter | -0.09 | -0.13 – -0.06 | **<0.001** |
| Error in exponent parameter (1<sup>st</sup> order) | -0.04 | -0.08 – -0.01 | **0.006** |
| Error in exponent parameter (2<sup>nd</sup> order) | 0.15 | 0.12 – 0.18 | **<0.001** |
| Observations | 4688 |  |  |
| R<sup>2</sup> / R<sup>2</sup> adjusted | 0.031 / 0.030 |  |  |

**Figure S1.**
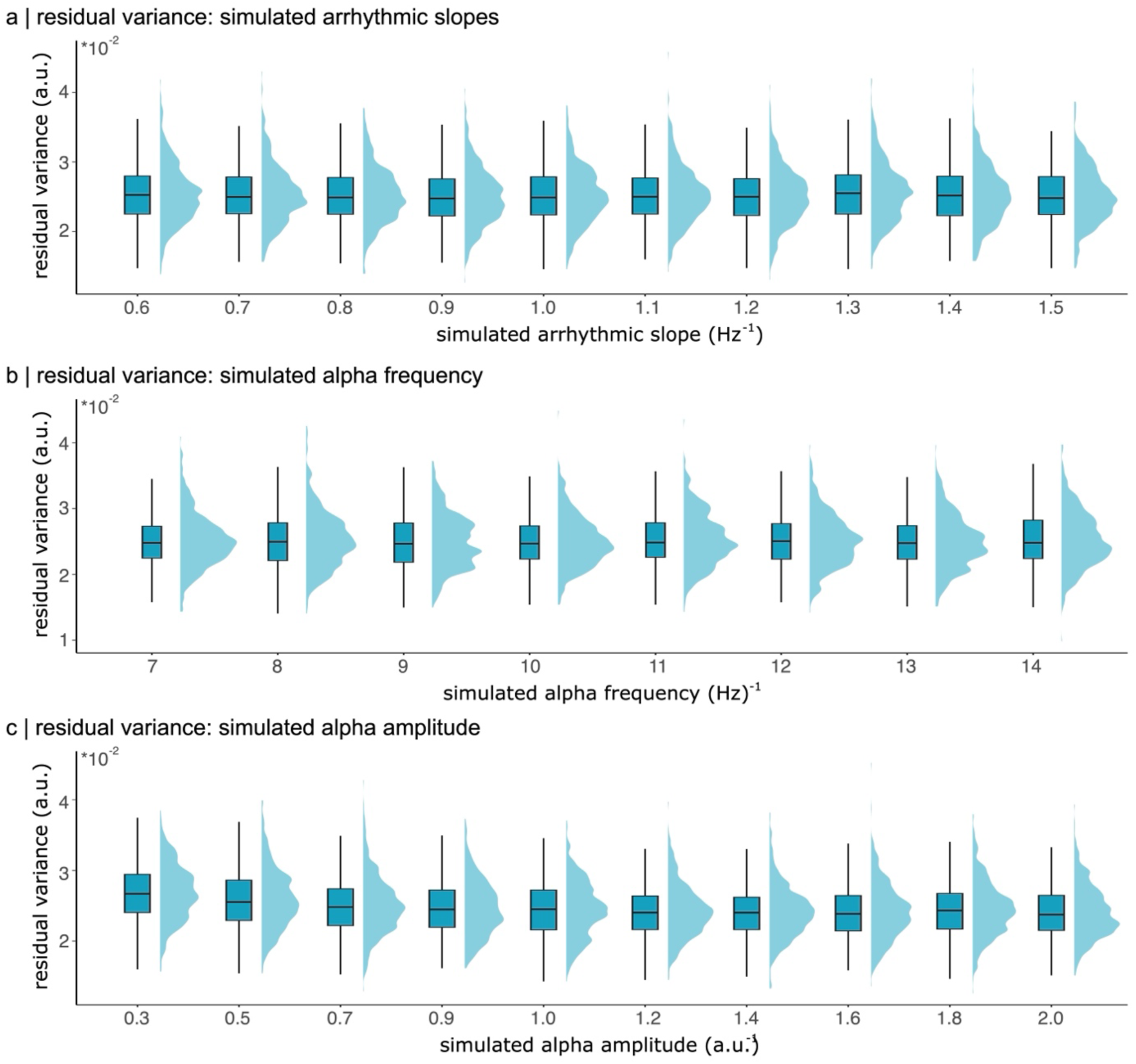
Residual variance of modelled spectra for all simulations. Box plots of the residual model variance (i.e., mean squared error) for all simulations with no relationship between rhythmic alpha power and the arrhythmic exponent. We systematically simulated neural time series data to parameterize with varying levels of (a) arrhythmic exponent, (b) alpha centre frequencies, and (c) peak alpha amplitude. The model fit of *ms-specparam* did not systematically vary as a function of the simulated parameters.

**Figure S2.**
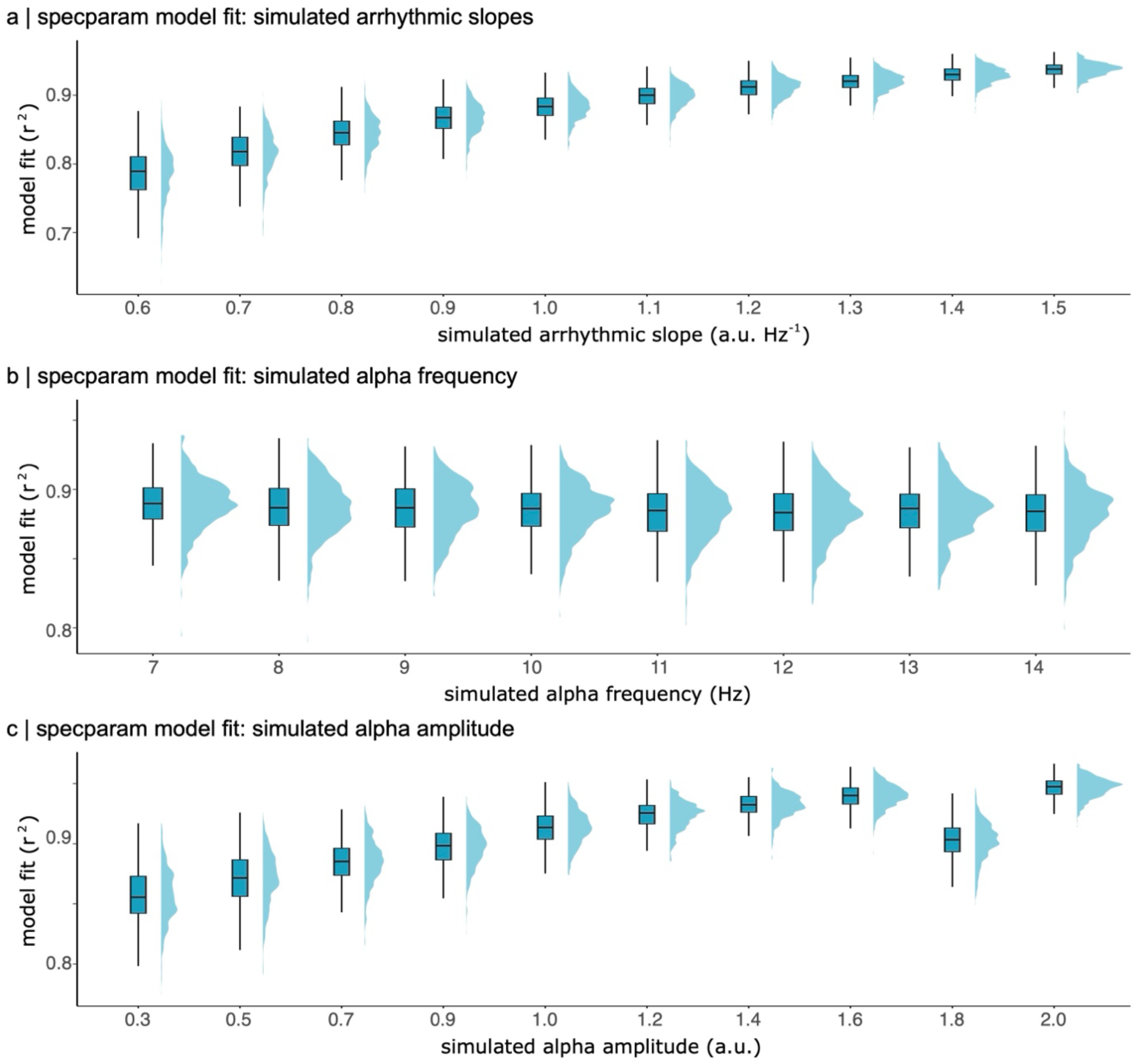
Specparam model goodness of fit metrics for all simulations. Box plots of the goodness of fit (r^2^) for the specparam model for all simulations with no ground-truth relationship between rhythmic alpha power and the arrhythmic exponent (i.e., Figure 2). We systematically simulated neural time series data to parameterize with varying levels of (a) arrhythmic exponent, (b) alpha centre frequencies, and (c) peak alpha amplitude. The goodness of fit of *ms-specparam* varied systematically as a function of modelled arrhythmic slope and simulated alpha amplitude. The alpha amplitude effect can be explained as larger amplitude alpha peaks by definition have higher SNR. The arrhythmic exponent effects can be attributed to a flatter spectral slope lowering the total sum of squares, and therefore the overall r^2^. These relationship between spectral parameters and goodness of fit metrics have been documented previously (Donoghue et al., 2020; L. E. Wilson et al., 2024). However, as shown in Figure S1, there is no discernible effect of simulated model parameters on the residual variance left unexplained by the model.

**Figure S3.**
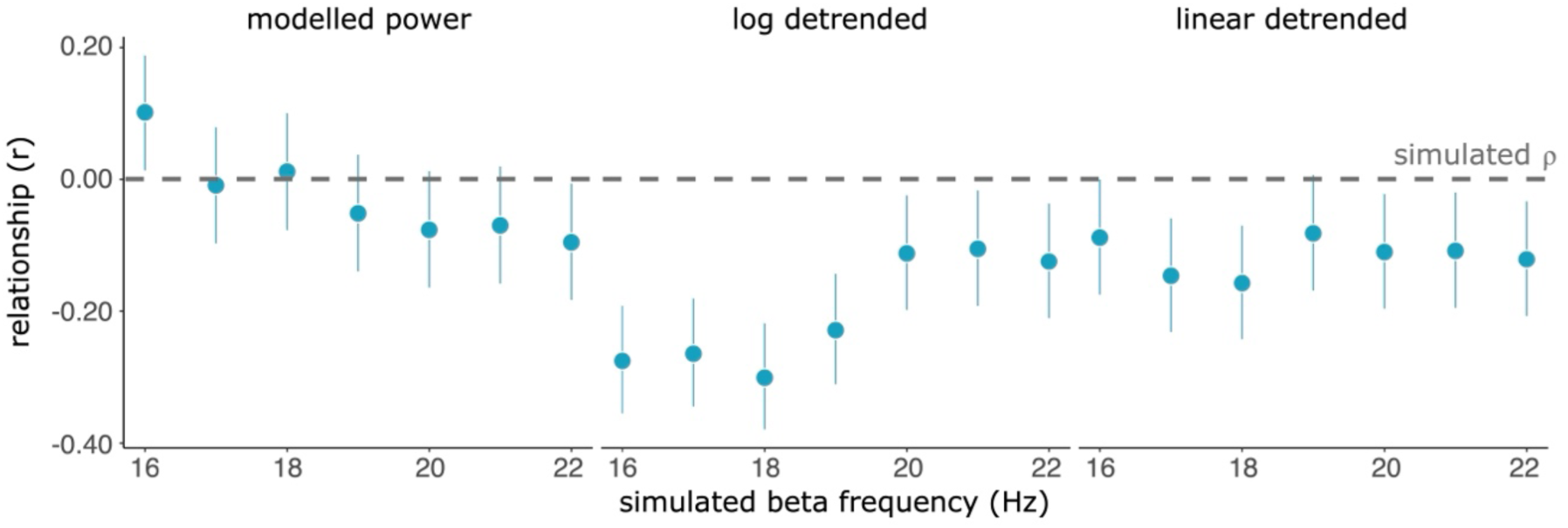
Spurious correlations between log detrended beta power and exponent. To verify that the observed spurious relationships between rhythmic power and arrhythmic exponent are not specific to alpha rhythms, we simulated synthetic neural time series data and systematically varied beta peak centre frequency. Similar to the alpha peak centre frequency findings, modelled beta amplitudes were closer to the ground-truth relationship than log- and linear- detrended rhythmic beta power. Rhythmic beta power was weakly correlated to the arrhythmic exponent when computed using detrending methodology. This effect was most evident for low-beta (> 20 Hz).

**Figure S4.**
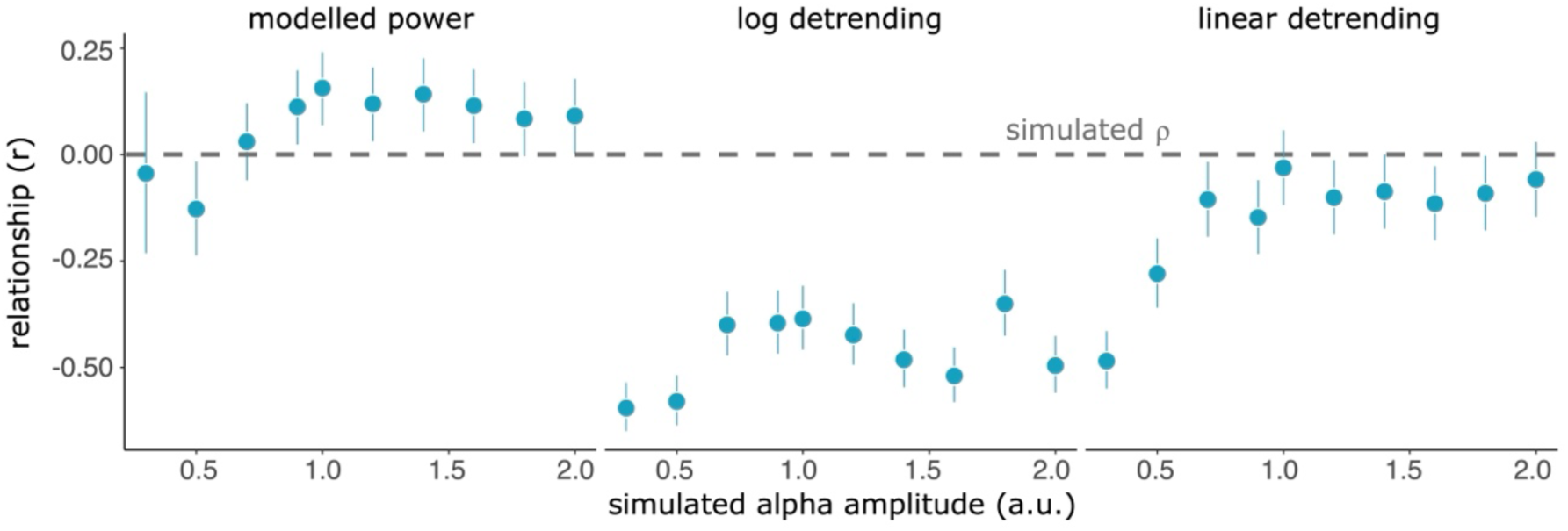
Spurious correlations between log detrended alpha power and exponent for different values of simulated amplitudes. Scatter plot of the relationship between rhythmic alpha power and arrhythmic exponent for various values of simulated rhythmic alpha amplitude. Just as the results presented in the main text, modelled power computed as the maximum amplitude of modelled Gaussian peaks is the closest to the ground truth (i.e., ⍴ = 0, dashed grey line). In contrast, log- and linear- detrending of the power spectrum introduces systematic spurious correlations between estimated rhythmic alpha power and the arrhythmic exponent. This is most evident for the lowest amplitude peaks (peak amplitude < 0.7).

**Figure S5:**
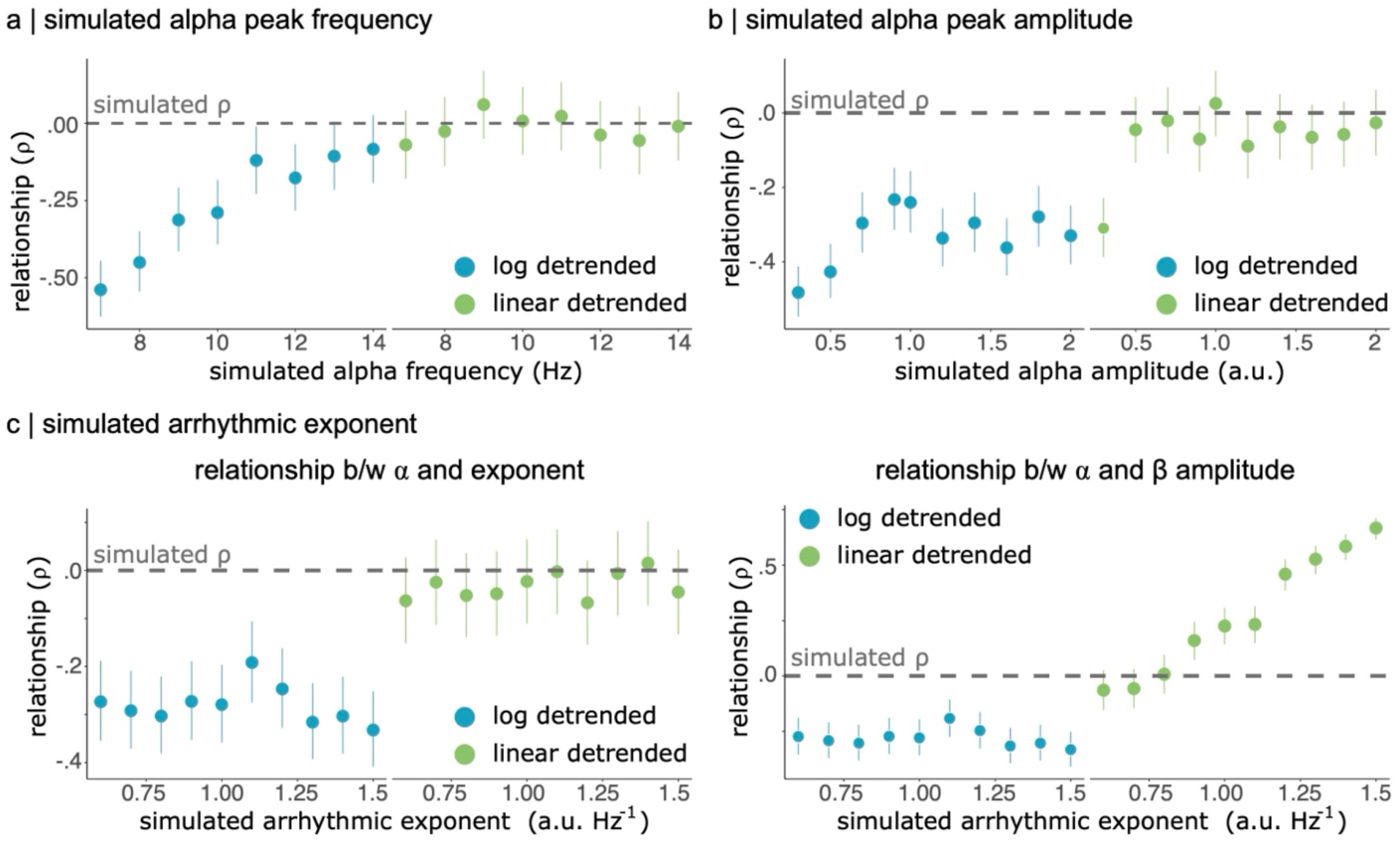
Maximum alpha amplitude of detrended spectrum induces spurious correlations between spectral model outputs. We estimated the correlation between rhythmic alpha power and arrhythmic exponent across three different sets of simulations (a, b, & c left panel) and the relationship between alpha and beta power for various values of simulated arrhythmic exponent. The ground truth correlation is zero, indicated by the dotted line for all these simulation experiments. Log- and linear- detrended power were defined as the maximum power within the alpha range after detrending the power spectrum in log or linear space. This is in contrast to taking the mean within the pre-defined alpha band. (a) Simulated alpha center frequencies. Spurious negative correlations of large magnitude (⍴ < -0.40) between alpha power and arrhythmic exponent were observed for log-detrended power. This is especially evident when the center frequency of the alpha oscillation lies at the edge of the narrow-band definition (i.e., 7-8 Hz). (b) Simulated alpha peak amplitudes. Log-detrended power induced large spurious correlations. (c) Left panel: Simulated arrhythmic exponents. Log-detrended power induced large spurious correlations. Right panel: Scatter plot of the relationship between rhythmic alpha and beta power for various simulated arrhythmic exponents. While synthetic time series data were simulated to have no linear relationship between alpha and beta amplitudes, spurious correlations were observed between alpha and beta band power for log- and linear-detrending methods. Error bars represent 95% CI.

**Figure S6.**
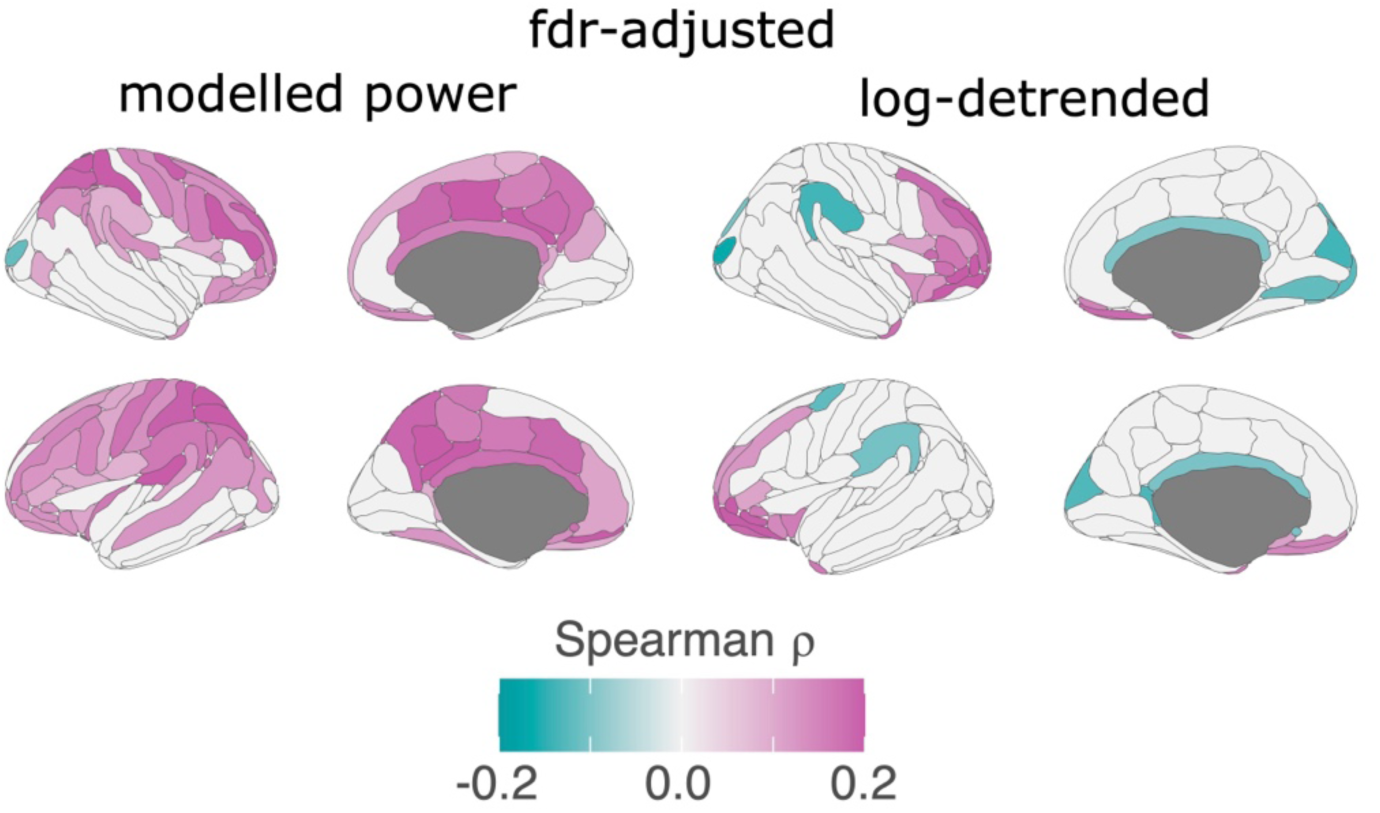
FDR-adjusted topographic map of the relationship between alpha power and exponent. The relationship between arrhythmic exponent and rhythmic alpha power for each parcel of the Desterieux atlas modelled (left) and log-detrended power (right). The topographic maps are thresholded (*p_FDR_* < 0.05) and depict significant relationships after correcting for multiple comparisons. While there are significant relationships between arrhythmic exponent and rhythmic alpha amplitude for both methods, the directionality and spatial extent of the relationship greatly depend on the method of quantifying rhythmic power.

**Figure S7.**
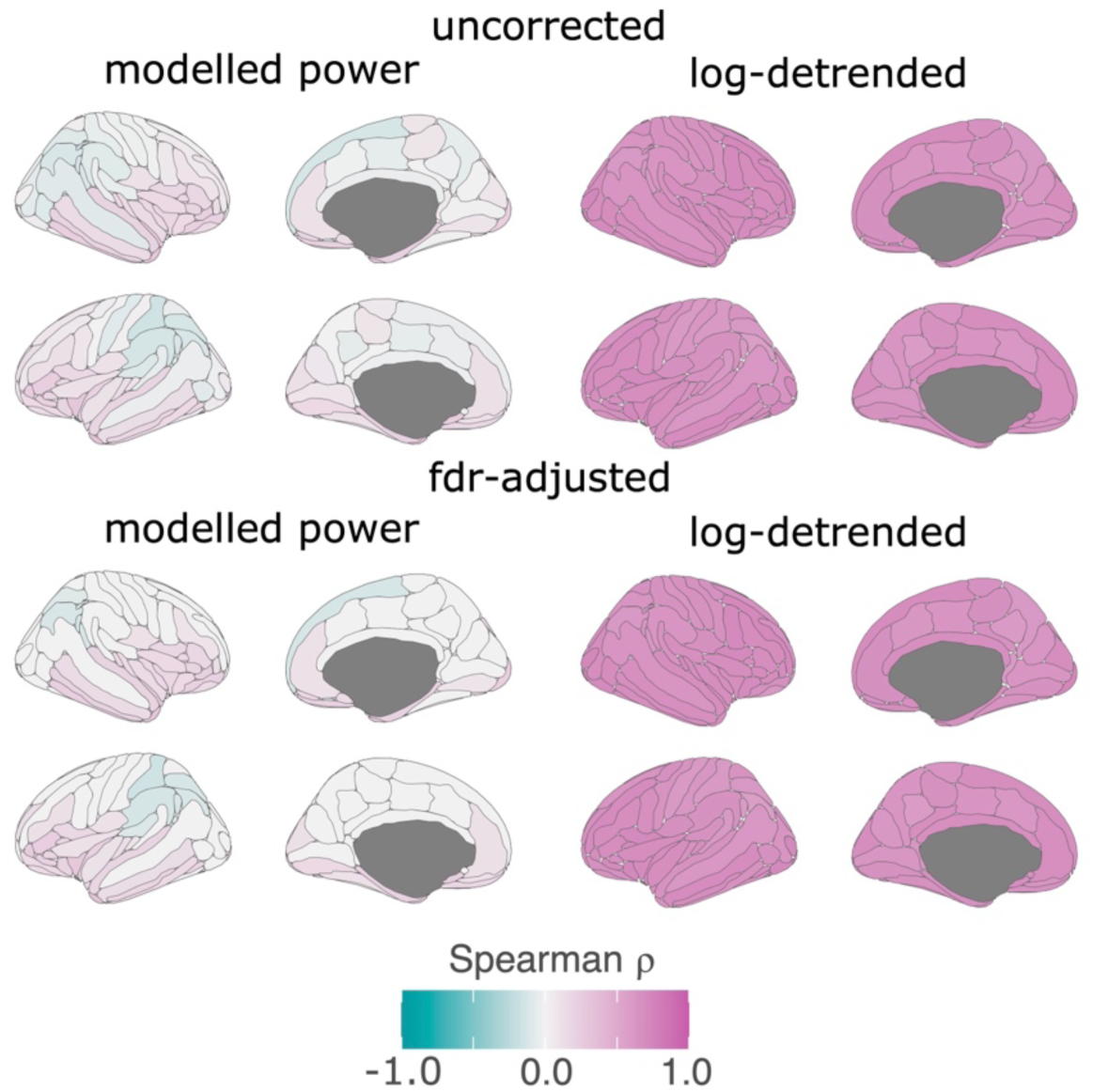
Topographic maps of the relationship between resting-state rhythmic alpha and beta power. Topographic maps depicting the relationship between rhythmic alpha and beta power for each parcel of the Desterieux atlas based on modelled (left) and log-detrended power (right). The unthresholded (top panel) and thresholded (*p_FDR_* < 0.05, bottom panel) topographic maps underscore the dramatic differences between both methodologies. Log-detrended power estimates (right panel) induce large correlations between the estimates of rhythmic alpha and beta power due to residual contributions of the arrhythmic exponent. In contrast, rhythmic alpha and beta power are weakly correlated to one another when relying on estimates of modelled power (left panel).

**Figure S8.**
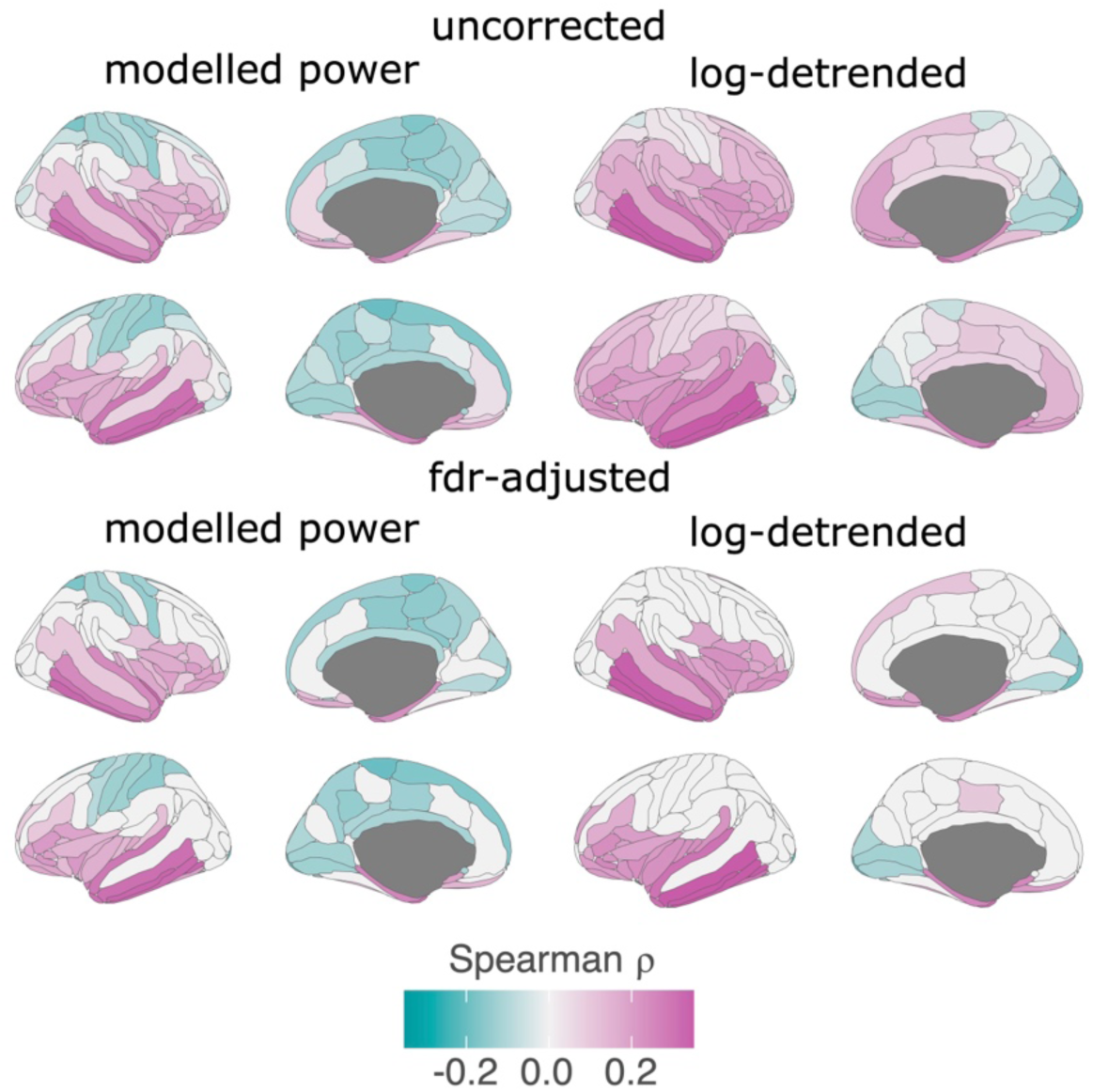
The relationship between resting-state alpha power and age diverges between modelling and detrending methods. The relationship between rhythmic alpha power and chronological age for each parcel of the Desterieux atlas modelled (left) and log-detrended power (right). The unthresholded (top panel) and thresholded (*p_FDR_* < 0.05, bottom panel) topographic maps demonstrate the differences in interpretation when utilizing different methods of quantifying rhythmic power. While age generally positively relates to log-detrended alpha power, this relationship inverts for modelled alpha power. We attribute these differences in findings to the arrhythmic exponent which is similarly altered by aging.

**Figure S9.**
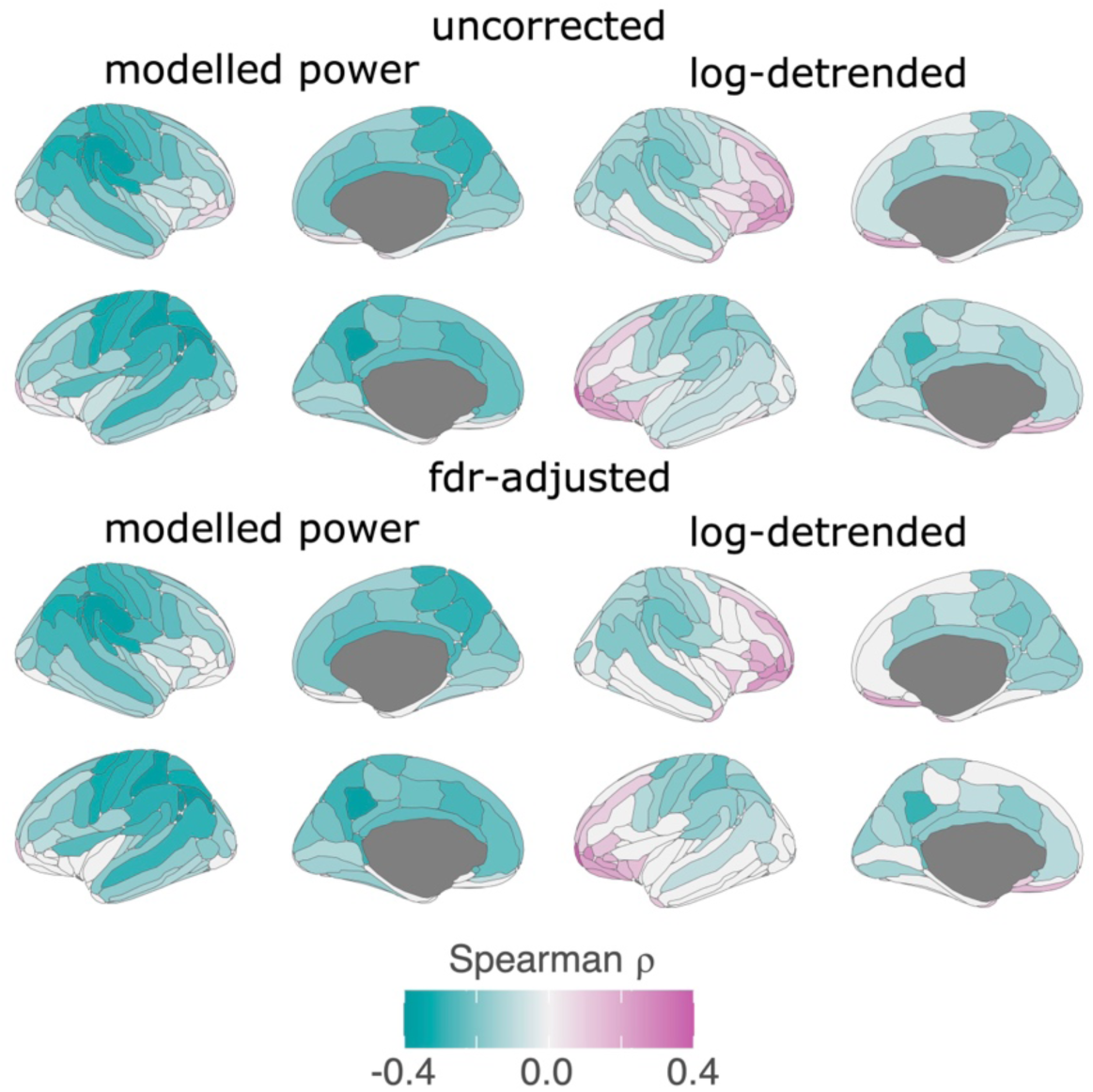
The relationship between resting-state beta power and arrhythmic exponent diverges between modelling and detrending methods. The relationship between rhythmic beta power and arrhythmic exponent for each parcel of the Desterieux atlas modelled (left) and log-detrended power (right). The unthresholded (top panel) and thresholded (*p_FDR_* < 0.05, bottom panel) topographic maps demonstrate the differences in interpretation when utilizing different methods of quantifying rhythmic power. While the arrhythmic exponent relates positively to frontal log-detrended beta power, this relationship inverts for modelled beta power.

